# Bacterial cGAS senses a viral RNA to initiate immunity

**DOI:** 10.1101/2023.03.07.531596

**Authors:** Dalton V. Banh, Cameron G. Roberts, Adrian Morales Amador, Sean F. Brady, Luciano A. Marraffini

## Abstract

CBASS immunity protects prokaryotes from viral (phage) attack through the production of cyclic dinucleotides which activate effector proteins that trigger the death of the infected host. How bacterial cyclases recognize phage infection is not known. Here we show that staphylococcal phages produce a highly structured 400-nt RNA, termed <u>C</u>BASS-<u>a</u>ctivating <u>b</u>acteriophage RNA (cabRNA), that binds to a positively charged surface of the CdnE03 cyclase and promotes the synthesis of the cyclic dinucleotide cGAMP. Phages that escape CBASS immunity harbor mutations that lead to the generation of a longer form of the cabRNA that cannot activate CdnE03. Since the mammalian cyclase OAS1 also binds viral dsRNA during the interferon response, our results reveal a conserved mechanism for the activation of innate antiviral defense pathways.

## INTRODUCTION

As a result of an evolutionary arms race, bacteria have evolved numerous immune strategies to counter infection by predatory viruses known as bacteriophages (or phages). Remarkably, many recently discovered antiviral systems in bacteria share structural and functional homology to components of metazoan innate immunity ^1, 2^. One key example of this ancestral connection includes <u>c</u>yclic oligonucleotide-<u>b</u>ased <u>a</u>ntiphage <u>s</u>ignaling <u>s</u>ystems (CBASS) in bacteria ^3, 4^, which are analogous to the <u>c</u>yclic <u>G</u>MP-<u>A</u>MP <u>s</u>ynthase (cGAS)-<u>st</u>imulator of <u>in</u>terferon <u>g</u>enes (STING) antiviral pathway in metazoans ^5, 6^. CBASS contain two core components: a cGAS/DncV-like <u>c</u>yclic <u>din</u>ucleotidyltransfer<u>ase</u> (CD-NTase, or Cdn) enzyme that generates cyclic nucleotides in response to phage infection ^7–9^, and an effector protein that binds the cyclic nucleotides to trigger the death or growth arrest of the host and thus inhibit viral propagation ^3, 4, 9^. In addition to the cyclase and effector genes, CBASS operons can encode for accessory proteins that are used for their classification into four major types ^4^. Type I CBASS comprise the minimal and most abundant architecture (42% of the analyzed operons have this composition). Type II CBASS (39%, the second most common) encode additional genes with ubiquitin-associated domains. Type III CBASS (10%) include regulatory genes that encode eukaryotic-like HORMA and TRIP13 domains ^10^. Finally, Type IV CBASS are the rarest and is enriched in archaea ^4^.

A central aspect of cyclic nucleotide-based immunity is the mechanism of activation of the cyclase; i.e., how the enzyme senses viral infection to begin the synthesis of the second messenger. For human cGAS, this is achieved through direct interaction with viral double-stranded DNA present in the cytosol ^6, 11–14^. Other cGAS homologs present in animals, however, can sense RNA instead of DNA ^15–18^. In contrast to metazoan cGAS-based immunity, the mechanisms that govern cyclase activation during the bacterial CBASS response are poorly understood. Biochemical analyses of bacterial cyclases from a variety of divergent CBASS operons demonstrated that some of these enzymes are constitutively active *in vitro* ^9^, suggesting that their activity *in vivo* is negatively regulated and only unleashed upon phage recognition. For instance, it has been proposed that the type II CBASS cyclase DncV from *Vibrio cholerae* is inhibited by folate-like molecules ^19^. These metabolites are presumably depleted during infection, a decrease that triggers second messenger production. However, there are also many examples of CBASS cyclases that are inactive *in vitro,* and therefore must require a mechanism of activation to initiate the immune response. This seems to be the case for the *E. coli* type III CBASS, which is activated *in vitro* through recognition of peptides (presumably of phage origin) by HORMA domain proteins that then form a complex with the cognate cyclase to initiate cyclic nucleotide synthesis ^10^. Interestingly, this system requires not only the HORMA domain protein, but also the binding of dsDNA by the cyclase for activation *in vitro* ^10^. How immunity is initiated by minimal CBASS operons that lack regulatory genes is not known. Here we investigated the mechanism of activation of the cyclase present in the minimal type I CBASS from *Staphylococcus schleiferi.* We found that both *in vitro* and *in vivo*, the binding of a structured RNA produced by staphylococcal phages during infection leads to the synthesis of cGAMP, which in turn activates a transmembrane effector to induce abortive infection.

## RESULTS

### CBASS protects staphylococci from phage infection

Bioinformatic analyses have previously uncovered more than 100 CBASS operons in diverse *Staphylococcus* strains ^4^, none of which, however, have been tested experimentally. We decided to characterize a type I-B CBASS present in *Staphylococcus schleiferi* strains 2142-05, 2317-03, and 5909-02 ^20^, hereafter designated Ssc-CBASS (Fig. S1A). This system consists of a two-gene operon harboring a Cdn belonging to the E clade, cluster 3 (Ssc-CdnE03) ^9^, and a transmembrane effector, Cap15, that was recently demonstrated to limit phage propagation by disrupting the host membrane ^21^. Since we were unable to find a phage that infects this organism, we cloned Ssc-CBASS, as well as the cyclase gene alone as a control, into the staphylococcal vector pC194 ^22^ for expression in the laboratory strain *Staphylococcus aureus* RN4220 ^23^. The resulting strain was infected with four lytic phages on soft-agar plates to enumerate plaque formation. We found that Ssc-CBASS, but not Ssc-CdnE03 alone, strongly reduced the propagation of Φ80ɑ-vir [a lytic derivative of the temperate phage Φ80ɑ ^24^ created for this study] and ΦNM1γ6 ^25^, but not for ΦNM4γ4 ^26^ (for which plaque size was reduced however) nor Φ12γ3 ^27^ (Fig. 1A). Similar results were obtained using a chromosomally expressed Ssc-CBASS (Fig. S1B). Consistent with previous reports ^3^, infection of liquid cultures with Φ80ɑ-vir at different multiplicity of infection (MOI) showed that Ssc-CBASS confers robust immunity and enables a complete recovery of the bacterial population at low phage concentrations (Fig. 1B). In addition, enumeration of colony-forming units (CFU) immediately before and after phage infection at MOI 5 indicated that Φ80ɑ-vir (Fig. 1C), but not the Ssc-CBASS-insensitive ΦNM4γ4 phage (Fig. S1C), causes loss of cell viability. This initial reduction is followed by an increase in CFU (presumably due to the growth of uninfected cells) that reflects Ssc-CBASS-mediated immunity against Φ80ɑ-vir. This was not observed after infection with ΦNM4γ4, where the CFU count decreased with time. As expected, plaque-forming units (PFU) enumerated in these samples were in line with the CFU counts, demonstrating the inability of Φ80ɑ-vir to detectably propagate (Fig. 1D) compared to a steady increase in ΦNM4γ4 PFUs over time (Fig. S1D). Altogether these data show that, similarly to other species, CBASS defense protects staphylococcal populations by preventing the growth of infected hosts to limit viral propagation.

**Figure 1.**
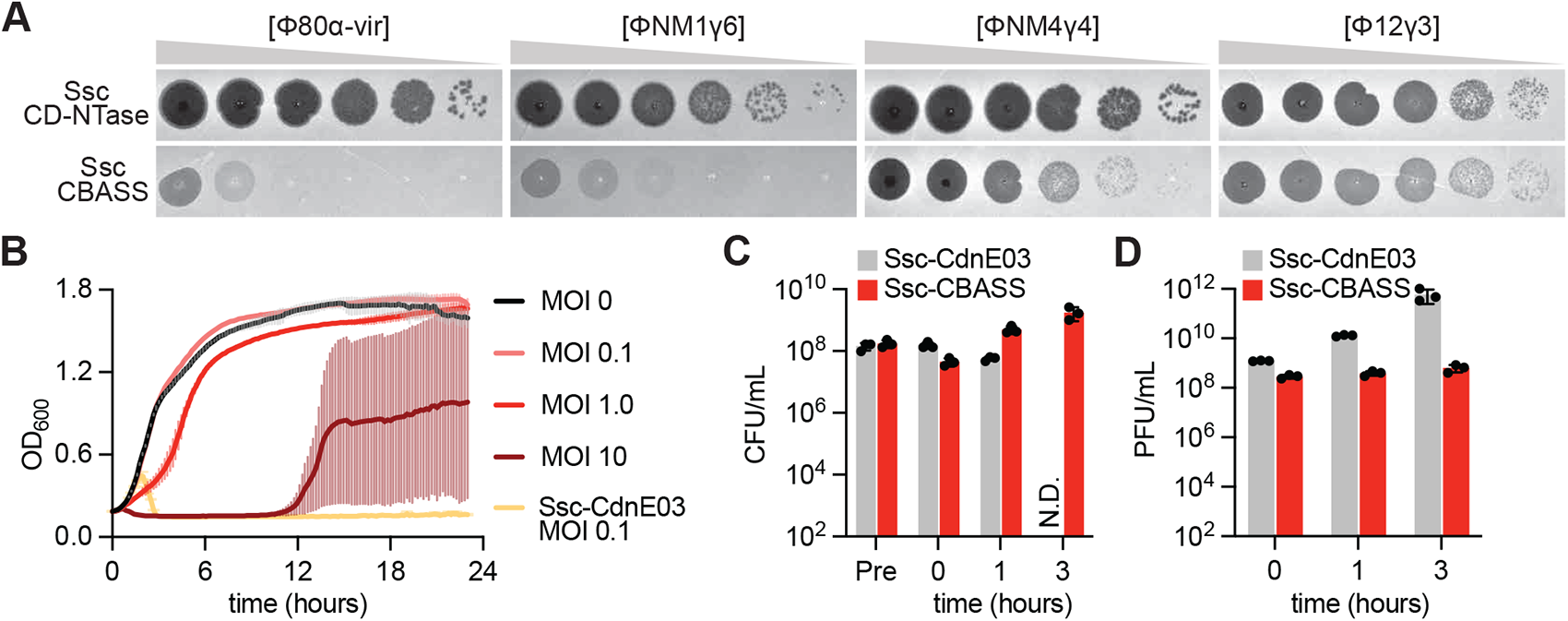
CBASS confers anti-bacteriophage defense in staphylococci via abortive infection. **(A)** Detection of phage propagation after spotting ten-fold dilutions of the lytic DNA phages, Φ80α-vir, ΦNM1γ6, ΦNM4γ4, and Φ12γ3 onto lawns of *S. aureus* RN4220 harboring a plasmid-borne either incomplete (Ssc-CdnE03 alone) or intact Ssc-CBASS operon. **(B)** Growth of staphylococci harboring either an incomplete (Ssc-CdnE03 alone) or intact Ssc-CBASS operon measured by optical density at 600 nm after the addition of Φ80α-vir at a multiplicity of infection (MOI) of 0, 0.1, 1, or 10. The mean of three biological replicates ± SD is reported. **(C)** Enumeration of colony-forming units (CFU) from cultures harboring Ssc-CdnE03 alone or Ssc-CBASS immediately before infection (Pre), after initial absorption of the phage (0 h), after one lytic cycle (1 h), and after complete culture lysis (3 h) by Φ80α-vir at MOI 5. Mean ± SEM of three biological replicates is reported. **(D)** Same as **(C)** but enumerating of plaque-forming units (PFU). Mean ± SEM of three biological replicates is reported.

### A 400-nucleotide phage RNA binds and activates Ssc-CdnE03

Next, we investigated how Ssc-CBASS is activated by Φ80ɑ-vir. We first considered the possibility of transcriptional activation of the operon during infection, as was reported for the type III CBASS of *Escherichia coli* upec-117 ^28^. RT-qPCR, however, failed to detect an increase in the transcription of the Ssc-CBASS genes upon infection (Fig. S2A). In addition, overexpression of the full operon, or either the cyclase or the effector alone, did not result in cell toxicity in the absence of phage (Fig. S2B). Therefore, we decided to look for CBASS activators by performing nucleotide synthesis assays *in vitro* using purified Ssc-CdnE03. We incubated the cyclase with trace ^32^P-labeled NTPs and an excess of unlabeled NTPs and, following phosphatase treatment, we visualized the reaction products using thin-layer chromatography ^9^. We tested *S. aureus* RN4220 crude lysate, purified Φ80ɑ-vir particles, host genomic DNA, phage genomic DNA, and total RNA from both uninfected and infected *S. aureus* RN4220 cells. Strikingly, only RNA isolated from cells infected with Φ80ɑ-vir enabled the generation of a cyclic nucleotide product by wild-type Ssc-CdnE03 (Fig. 2A), but not the active site mutant D86A,D88A that fails to mediate immunity (Fig. S2C). To determine the cyclase product, we used different radiolabeled NTPs and found that ATP and GTP are both necessary and sufficient for product formation (Fig. S2D). Further analysis of this product by LC-MS defined it as an isomer (3’,3’ or 3’,2’) of cyclic guanosine monophosphate-adenosine monophosphate, cGAMP (Fig. S2E-F and Supplementary Text).

**Figure 2.**
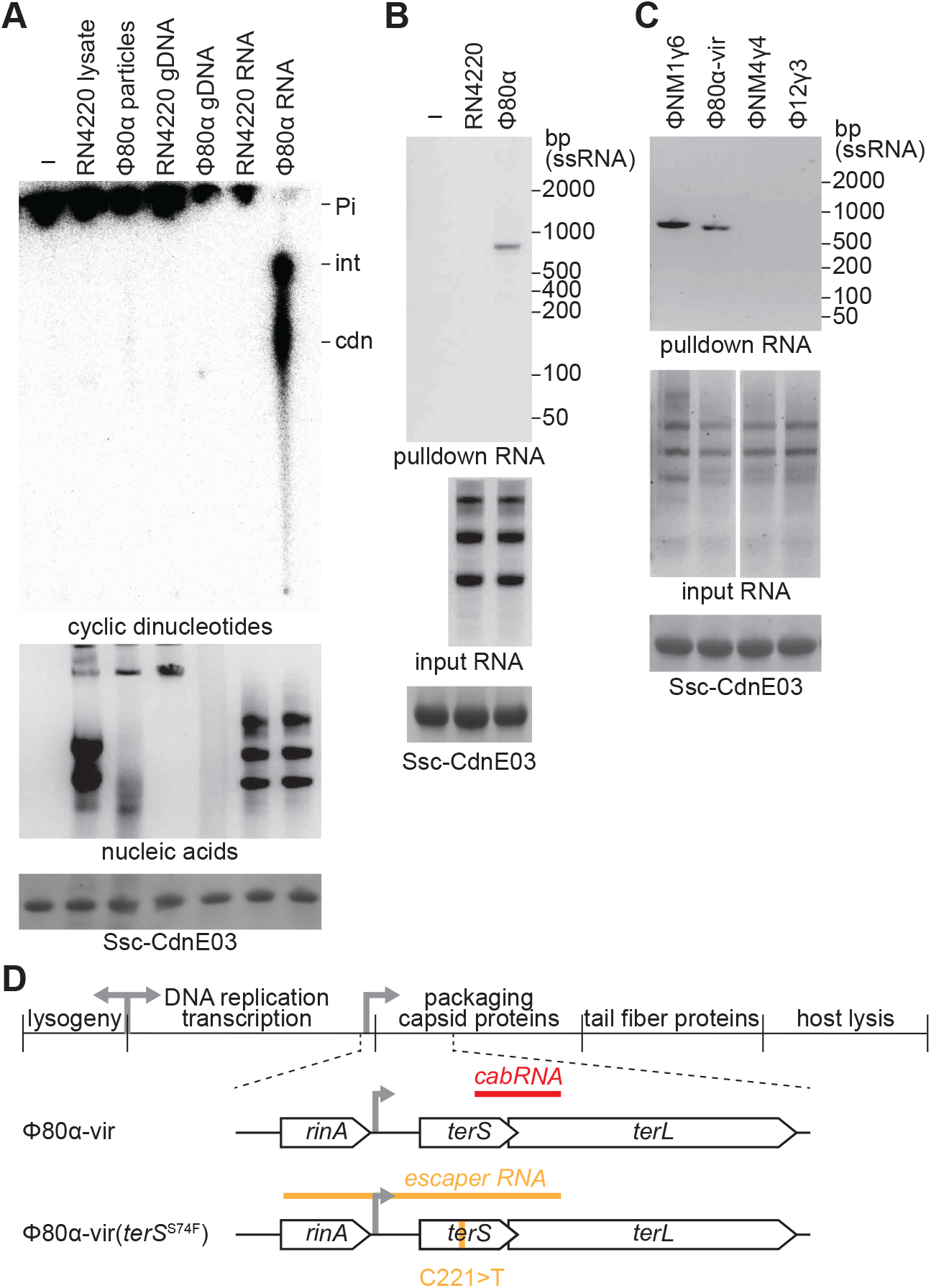
A viral RNA produced during infection activates Ssc-CdnE03 *in vitro*. **(A)** Thin-layer chromatography analysis of Ssc-CdnE03 products in the presence of the following: *S. aureus* RN4220 crude lysate, whole purified Φ80ɑ-vir particles, host genomic DNA (RN4220 gDNA), phage gDNA, and total RNA from *S. aureus* RN4220 in the presence or absence of Φ80ɑ-vir infection (before the completion of 1 lytic cycle). A representative image of multiple replicates is shown. An agarose gel stained with ethidium bromide (middle) and SDS-PAGE stained with Coomassie blue (bottom) are shown as loading controls. Pi, free phosphates; int, intermediate cyclase product; cdn, cyclic dinucleotide. **(B)** Agarose gel electrophoresis of the input and output RNA obtained after incubation of Ssc-CdnE03 with no RNA, total RNA extracted from uninfected staphylococci (RN4220) or from cells infected with Φ80ɑ-vir phage. An SDS-PAGE stained with Coomassie blue (bottom) is shown as a loading control. **(C)** Same as in **(B)**, but with input RNA extracted from staphylococci infected with ΦNM1γ6, Φ80α-vir, ΦNM4γ4, or Φ12γ3 phages. **(D)** Diagram of ϕ80α-vir and ϕ80α-vir(*terS*^S74F^) genomes with localization of the cabRNA and escaper RNA sequences, respectively. The location of the escaper mutation, C221>T, is shown.

To identify the activating RNA species, we purified a hexahistidyl-tagged, maltose-binding protein fusion of Ssc-CdnE03, and immobilized it to a cobalt resin column that was loaded with total RNA extracted from either infected or uninfected staphylococci. Extraction and separation of the nucleic acids bound by the cyclase revealed the presence of an RNA that migrated at approximately 800 nucleotides in length (compared to an ssRNA ladder) that was pulled down only from the RNA fraction of infected, but not uninfected cells (Fig. 2B). We repeated this assay with RNA obtained from cells infected with other phages and isolated a similar species for the Ssc-CBASS-sensitive ΦNM1γ6 phage, but not for the resistant ΦNM4γ4 and Φ12γ3 viruses (Fig. 2C). We subjected both isolated RNA species to next-generation sequencing to determine their origin. For the Φ80ɑ-vir RNA bound to the cyclase we found that reads mapped to a 400-nucleotide region beginning within the *gp40* gene and extending into *gp41*, which encode the terminase small and large subunits, TerS and TerL, respectively (Fig. 2D and Supplementary Sequences). Similar results were obtained for the cyclase-bound RNA generated during ΦNM1γ6 infection (Fig. S2G and Supplementary Sequences). We named this viral-derived RNA the “<u>C</u>BASS-<u>a</u>ctivating <u>b</u>acteriophage RNA” (cabRNA). Interestingly, we also detected reads for a 400-nucleotide host RNA derived from *addB*, which encodes one of the subunits of the AddAB helicase/nuclease complex involved in homologous-directed DNA repair ^29^ (Fig. S2H and Supplementary Sequences). This RNA was not detected in the material pulled-down by the cyclase after incubation with total RNA from the host, in the absence of phage infection, using RT-PCR. This result suggests that, as is the case for the cabRNA, the host RNA associated with Ssc-CdnE03 is generated during the Φ80ɑ-vir lytic cycle. Finally, we purified the RNAs obtained during the pull-down assays and found that they activate cGAMP production *in vitro* (Fig. S2I).

### Secondary structures within the cabRNA are required to activate Ssc-CdnE03

The electrophoretic migration of the cabRNA, higher than its nucleotide length (runs similarly to a ∼800-nt ssRNA, but is actually 400-nt long), suggests the existence of secondary structures that may be important for cyclase activation. Using ViennaRNA software ^30^ to predict such structures, we found several hairpins and double-stranded RNA (dsRNA) regions within the cabRNA (Fig. S3A). To test for the presence of these structures in the species pulled down by the cyclase, we used RNases T1 and III, which cleave ssRNA and dsRNA, respectively ^31^. We found that RNase III completely degraded the cabRNA, while RNase T1 did not affect this RNA (Fig. 3A). We also treated the total RNA extracted from infected staphylococci with these RNases as well as with another that degrades ssRNA, RNase A ^31^, and assayed for the ability of the treated samples to induce cGAMP production by Ssc-CdnE03 (Fig. 3B). While both RNases T1 and A degraded most of the RNA extracted from infected cells, the treated samples were still able to activate the cyclase. In contrast, RNase III treatment showed a limited impact on the degradation of the total RNA but completely abrogated the ability of the total RNA to induce cGAMP production. These results demonstrate that dsRNAs, but not ssRNAs, produced during Φ80ɑ-vir infection are important for Ssc-CdnE03 activation.

**Figure 3.**
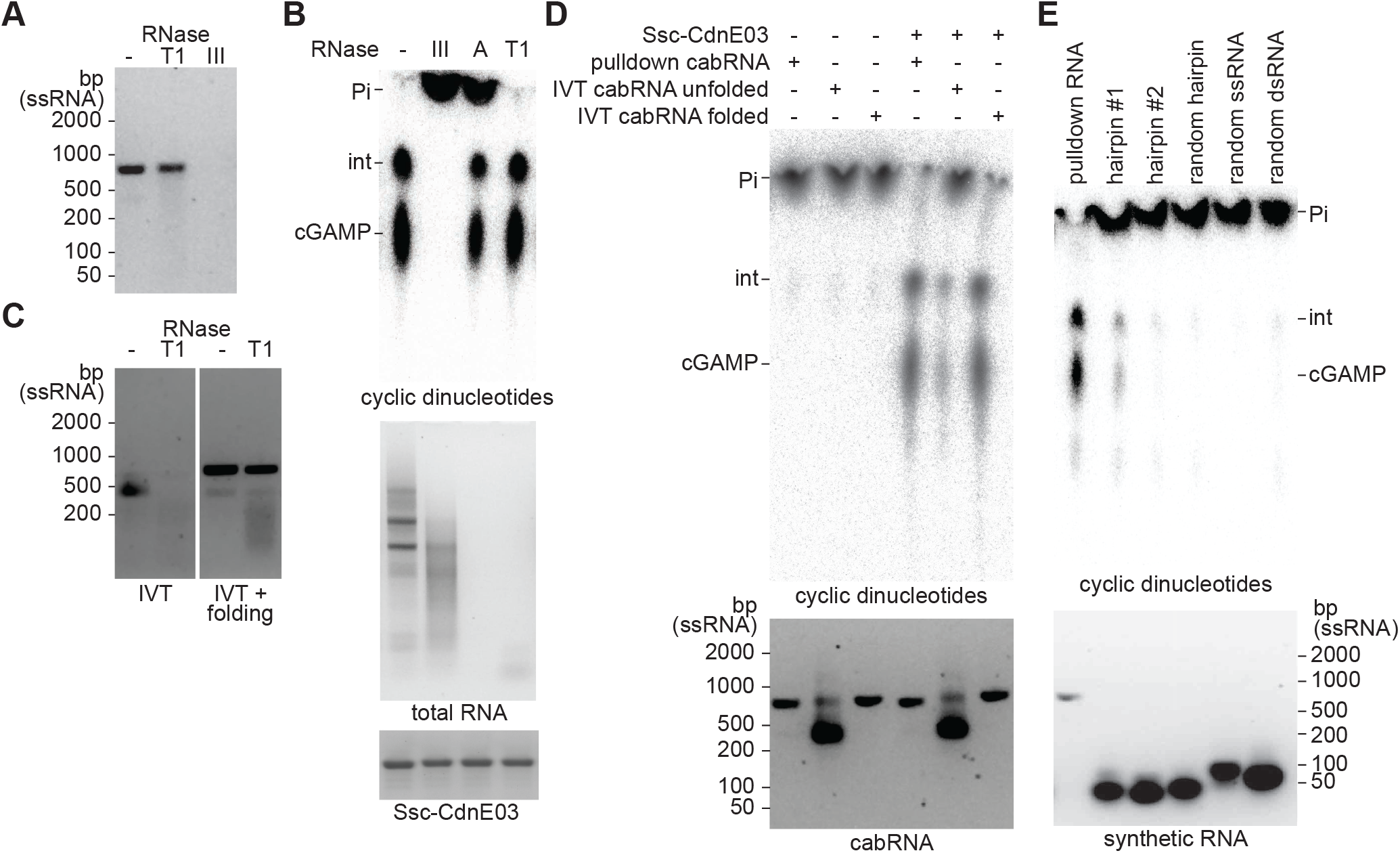
Secondary structures within the cabRNA are required for Ssc-CdnE03 activation. **(A)** Agarose gel electrophoresis of pulled-down cabRNA treated with RNases T1 or III. **(B)** Thin-layer chromatography analysis of Ssc-CdnE03 products in the presence of total RNA extracted from infected cells and treated with RNases III, A, T1, or untreated. A representative image of multiple replicates is shown. An agarose gel stained with ethidium bromide (middle) and SDS-PAGE stained with Coomassie blue (bottom) are shown as loading controls. Pi, free phosphates; int, intermediate cyclase product. **(C)** Agarose gel electrophoresis of *in vitro* transcribed (IVT) cabRNA, unfolded (left) and folded (right), untreated or treated with RNase T1. **(D)** Same as in **(B)** but incubating the cyclase with pulled-down, unfolded or folded IVT cabRNA. **(E)** Same as in **(B)** but incubating the cyclase with pulled-down RNA or different synthetic RNA oligonucleotides.

To determine if the cabRNA alone is sufficient for the activation of the cyclase, in the absence of other RNAs generated during infection, as well as to test the importance of RNA folding for this activation, we produced cabRNA *in vitro* using T7 RNA polymerase. The obtained transcript migrated similarly to a ssRNA of approximately 400 nucleotides after agarose gel electrophoresis and was completely digested by RNase T1 (Fig. 3C). This observation suggests that this *in vitro*-transcribed (IVT) cabRNA species lacks most of the secondary structures present in the cabRNA produced during infection. Therefore, we promoted the folding of the IVT cabRNA by heating it to 95°C for 5 minutes before slowly cooling the sample to room temperature. This treatment led to the generation of a cabRNA species with similar properties to that isolated from infected cells; i.e. migrated at ∼ 800 nt and was resistant to RNase T1 degradation (Fig. 3C). We tested the ability of the unfolded and folded IVT cabRNA to activate the cyclase and found that whereas the folded species induced cGAMP synthesis to the same levels of the cabRNA produced *in vivo*, the unfolded species triggered the production of substantially lower quantities of the cyclic dinucleotide, which we attribute to a low level of spontaneous folding of the IVT cabRNA in the sample used for this assay (Fig. 3D). We also tested the activating properties of an IVT RNA with a sequence complementary to that of the cabRNA (transcribed using the same template DNA but in the opposite direction), as well as the 400-nucleotide host RNA (transcribed from *addB*) that was found bound to the cyclase and its complementary RNA, in all cases heated and cooled to promote folding. None of these RNAs were found to activate Ssc-CdnE03 (Fig. S3A). Finally, we used synthetic RNA oligonucleotides with the sequences of the two most prominent predicted hairpins (hairpin-1 and -2, Fig. S3A) to determine their importance for cyclase activation. We found that, although at lower levels than the full cabRNA, hairpin-1, but not hairpin-2, induced cGAMP production (Fig. 3E). Other synthetic RNAs, including an unrelated hairpin structure, a dsRNA, and a ssRNA oligo (all with a similar size to hairpin-1 but with a different sequence, Table S5), failed to activate the cyclase. Altogether, these data indicate that specific secondary and/or tertiary structures within the cabRNA are essential for activation of Ssc-CdnE03.

We also investigated the ability of the cabRNA to activate Ssc-CBASS *in vivo*, in the absence of phage infection. To do this, we cloned the corresponding DNA sequence under the transcriptional control of an anhydrotetracycline (aTc)-inducible promoter on a staphylococcal expression vector ^32^. Given the abortive infection mechanism detected after Φ80ɑ-vir infection (Fig. 1B-D), we expected that, upon addition of aTc, induction of cGAMP synthesis by Ssc-CdnE03 and the subsequent activation of the membrane-disrupting effector would result in a proliferation defect and/or death of staphylococci. However, cabRNA transcription did not affect the growth of the cultures (Fig. S3C). To verify the plasmid-based expression of the cabRNA, we performed a pull-down assay using total RNA extracted from this strain after addition of aTc (Fig. S3D). Interestingly, most of the recovered cabRNA displayed an electrophoretic mobility consistent with the RNase T1-sensitive, unfolded form of the cyclase inducer (lower band). This observation suggests that the unfolded cabRNA can interact with, but not activate, Ssc-CdnE03. Given that mostly RNase III-sensitive, folded cabRNA is detected during pull-down assays using total RNA extracted from infected cells (Figs. 2B-C and S3D), we speculate that Φ80ɑ-vir infection is critical for the proper generation, modification, and/or folding of the inducer RNA.

### A conserved, positively charged surface within Ssc-CdnE03 binds the cabRNA

To investigate how the cabRNA interacts with Ssc-CdnE03, we obtained a structure prediction using AlphaFold (Fig. 4A and S4A). Similar to other characterized CD-NTases, Ssc-CdnE03 shares structural features and organization with mammalian OAS1 and cGAS despite low sequence similarity (∼20%) ^33^ (Fig. S4B). In particular, Ssc-CdnE03 and OAS1 share several core features: i) the common DNA polymerase β-like nucleotidyltransferase superfamily protein fold and conserved active site architecture, ii) a pocket on the backside of the active site with positive charge, iii) two positively charged residues (Arg and Lys) at the first helix of the PβCD domain, and iv) a surface exposed lysine and arginine along the enzyme “spine”. (Fig. S4C). Importantly, these features are associated with the sensing of dsRNA for OAS1 ^1, 6, 18^. To test if the positively charged surface of Ssc-CdnE03 is involved in cabRNA sensing, we substituted lysine residues present on this surface (K9, K13 Figs. 4A and S4C) for glutamic acid residues and assayed for cGAMP production *in vitro*. We found that the K9E and K13E substitutions substantially impaired the production of cGAMP in vitro and in vivo (Fig. 4B). To determine the role of this basic surface in cabRNA binding, we performed electrophoretic mobility shift assays using increasing concentrations of enzyme and observed that the substitution of K9, more than the K13E, notably affected the interaction of the cyclase with its inducer (Fig. 4C). Consistent with these *in vitro* results, CBASS immunity against Φ80ɑ-vir was most severely abrogated by the K9E mutation, and mildly reduced in staphylococci carrying the K13E mutant cyclase (Fig. 4D). Altogether these results demonstrate that the cabRNA interacts with a positively charged surface present in Ssc-CdnE03 to activate cGAMP synthesis and initiate the staphylococcal CBASS response.

**Figure 4.**
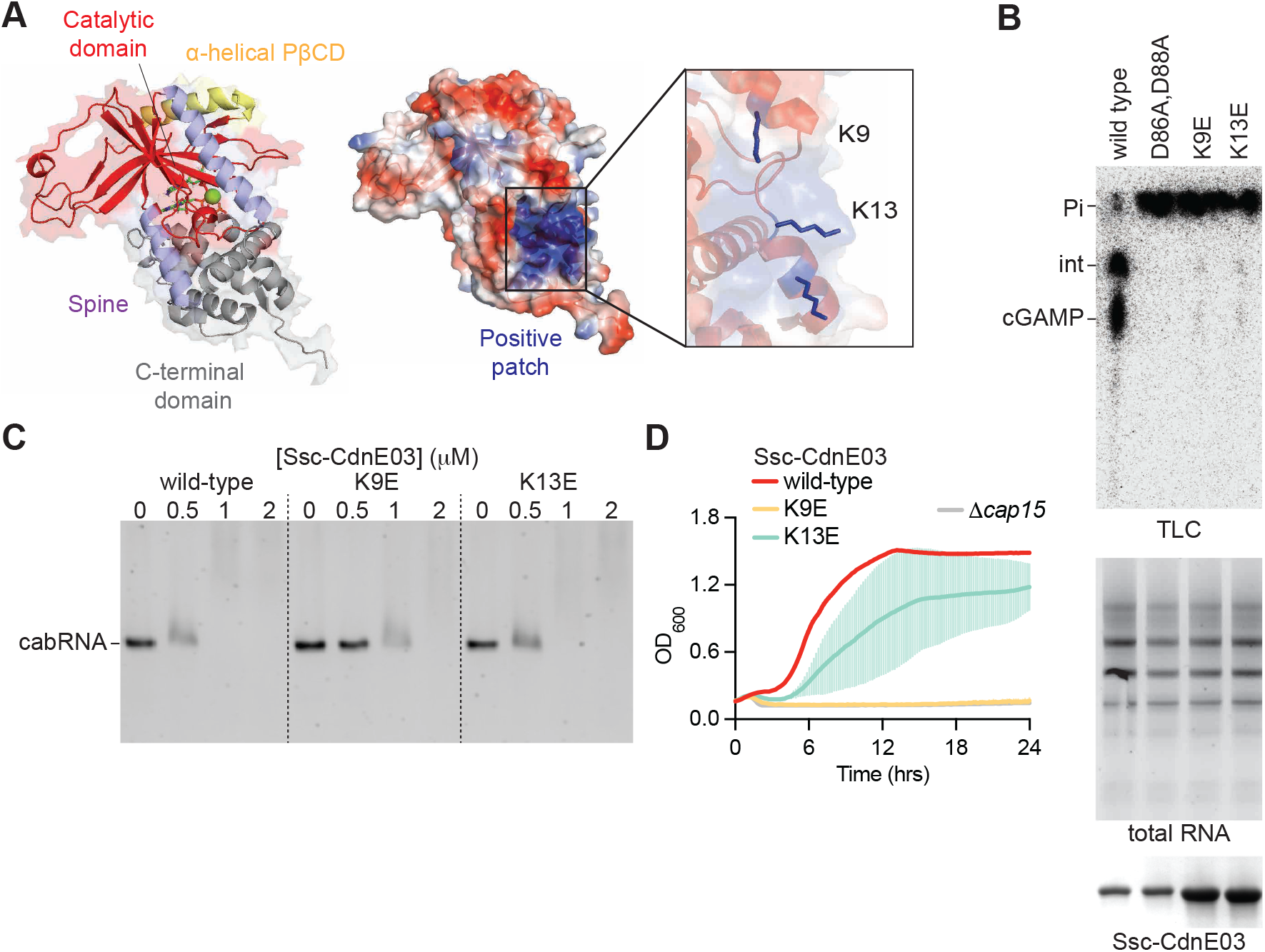
A positively charged surface within Ssc-CdnE03 binds the cabRNA to initiate immunity. **(A)** AlphaFold model of Ssc-CdnE03 displayed with surface electrostatics (-77 to 77, red to blue). Inset, positively charged region harboring the mutated lysine residues 9 and 13. **(B)** Thin-layer chromatography analysis of the products of different Ssc-CdnE03 mutants in the presence of total RNA extracted from infected cells. A representative image of multiple replicates is shown. An agarose gel stained with ethidium bromide (middle) and SDS-PAGE stained with Coomassie blue (bottom) are shown as loading controls. Pi, free phosphates; int, intermediate cyclase product. **(C)** Electrophoretic mobility shift assay of cabRNA in the presence of increasing concentrations of different Ssc-CdnE03 mutants. **(D)** Growth of staphylococci harboring the Ssc-CBASS operon with wild-type, K9E or K13E, Ssc-CdnE03 measured by optical density at 600 nm after infection with Φ80α-vir at an MOI of 10. The mean of three biological replicates ± SD is reported.

### Phage mutants that evade Ssc-CBASS immunity fail to produce cabRNA

To gain further insight into the mechanism of Ssc-CBASS induction, we sought to isolate phage mutants that can evade defense. Since we were unable to observe discrete Φ80ɑ-vir or ΦNM1γ6 plaques in our assays (Fig. 1A), something previously observed for defense mechanisms that mediate abortive infection ^34^, we used ethyl methanesulfonate (EMS) to introduce random mutations into a Φ80ɑ-vir population. Phages were plated on lawns of staphylococci harboring the Ssc-CBASS to isolate escape mutants that are able to form plaques (Fig. S5A). We then performed next-generation sequencing and detected Φ80ɑ-vir escapers carrying mutations (Fig. S5B). Many of these phages harbored nucleotide substitutions within the *gp46* gene, which encodes the scaffold protein for the viral capsid ^24^. This finding is consistent with recent reports of type II CBASS escape mutations in phage capsid genes ^35, 36^. Mutations generated missense amino acid substitutions in the scaffold protein and were corroborated to mediate the evasion of Ssc-CBASS immunity in infection and plaque formation assays; phenotypes that were reverted by the expression of wild-type Gp46 in the host (Fig. S5C-D). Next, we examined cabRNA production by these phages. Total RNA from staphylococci infected with phage expressing the *gp46*^E105D^ mutant or wild-type Φ80ɑ-vir mediated similar levels of cGAMP synthesis *in vitro* (Fig. S5E). In addition, pull-down assays using immobilized Ssc-CdnE03 retrieved the same cabRNA isolated from cells infected with wild-type viruses (Fig. 5D). Therefore, we conclude that these escapers that produce mutant capsids evade Ssc-CBASS immunity through a mechanism that does not affect the generation of cabRNA.

**Figure 5.**
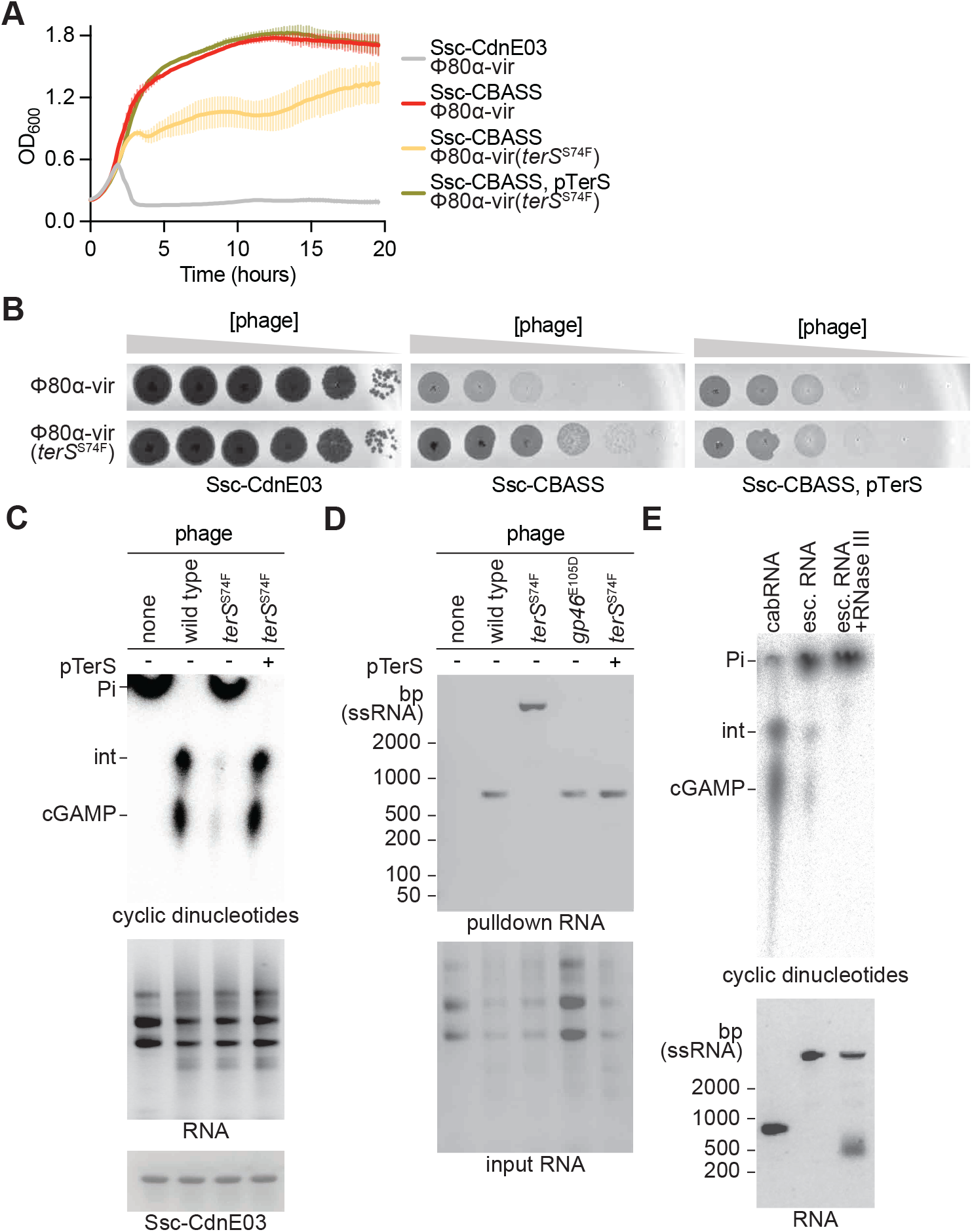
Phage mutants that evade Ssc-CBASS immunity fail to produce cabRNA. (A) Growth of staphylococci harboring either an incomplete (Ssc-CdnE03 alone) or intact Ssc-CBASS operon measured by optical density at 600 nm after infection with ϕ80α-vir or ϕ80α-vir(*terS*^S74F^) at MOI 1, the latter in the presence or absence of TerS overexpression using plasmid pTerS. The mean of three biological replicates ± SD is reported. **(B)** Detection of phage propagation after spotting ten-fold dilutions of Φ80α-vir or ϕ80α-vir(*terS*^S74F^) onto lawns of *S. aureus* RN4220 harboring either an incomplete (Ssc-CdnE03 alone) or intact Ssc-CBASS operon, the latter in the presence or absence of TerS overexpression using plasmid pTerS. **(C)** Thin-layer chromatography analysis of Ssc-CdnE03 products the presence of total RNA extracted from either uninfected staphylococci or cells infected with Φ80ɑ-vir or Φ80ɑ-vir(*terS*^S74F^), the latter in the presence or absence of TerS overexpression using plasmid pTerS. A representative image of multiple replicates is shown. An agarose gel stained with ethidium bromide (middle) and SDS-PAGE stained with Coomassie blue (bottom) are shown as loading controls. Pi, free phosphates; int, intermediate cyclase product. **(D)** Agarose gel electrophoresis of the input and output RNA obtained after incubation of Ssc-CdnE03 with total RNA extracted from uninfected staphylococci (RN4220) or from cells infected with Φ80ɑ-vir, Φ80ɑ-vir(*gp46*^E105D^) or Φ80ɑ-vir(*terS*^S74F^), the latter in the presence or absence of TerS overexpression using plasmid pTerS. **(E)** Same as **(C)** but in the presence of cabRNA, escaper RNA, or escaper RNA pre-treated with RNase III.

A different escaper mutation mapped to *terS*. This was a C to T transition that changes serine (UCU) 74 to phenylalanine (UUU) in this terminase subunit, located 6 nucleotides upstream of the cabRNA start. Infection of staphylococci harboring Ssc-CBASS with the Φ80ɑ-vir(*terS*^S74F^) mutant phage prevented bacterial growth (Fig. 5A), and resulted in the production of high numbers of viral particles (Fig. 5B). Total RNA from the infected cultures failed to stimulate cGAMP production (Fig. 5C). Surprisingly, pull-down assays revealed that the cyclase binds an RNA species generated during infection that is several hundreds of nucleotides larger than the cabRNA (Fig. 5D). RNA-seq of this species identified it as a 1,237-nt long transcript that starts at *gp39* [which encodes RinA, the transcriptional regulator of the late-expressed genes of Φ80ɑ-vir ^37^] and extends into *terL* (*gp41*), harbors the C to T mutation, and shares the same 3’ end with the cabRNA (Fig. 2D and Supplementary Sequences). Interestingly, in contrast to the cabRNA, which is resistant to degradation by RNase T1 and completely susceptible to RNase III (Fig. 3A), the 1,237-nt escaper RNA isolated from the pull-down assay was susceptible to RNase T1 and was only partially degraded by RNase III (Fig. S6A). This RNA retained a low level of cyclase activation, which was eliminated after RNase III treatment (Fig. 5E). On the other hand, a synthetic escaper RNA generated through *in vitro* transcription failed to induce cGAMP production (Fig. S6B). Altogether these results indicate that the “long” escaper RNA has a different secondary/tertiary structure than the cabRNA. This long RNA species mediates low levels of Ssc-CdnE03 binding and activation, but is not sufficient to trigger a full Ssc-CBASS response *in vivo*. Finally, we introduced the escape mutation into the *terS* homolog of ΦNM1γ6. We found that recombinant phages acquired resistance to Ssc-CBASS (Fig. S6C and Supplementary Sequences), a result that confirms that this *terS*^S74F^ mutation is sufficient to subvert Ssc-CBASS activation. Importantly, neither overexpression of wild-type TerS (Fig. S6D), Gp46 (Fig. S6E), nor the complete set of phage genes required for capsid formation (*gp40*-*gp47*, Fig. S6F), which expression was shown to form Φ80ɑ capsids in the absence of phage infection ^38^, impaired the growth of staphylococci carrying the Ssc-CBASS system, indicating that TerS, Gp46 or the full Φ80ɑ capsid, do not participate directly in Ssc-CdnE03 activation.

We also performed experiments to restore Ssc-CBASS immunity against Φ80ɑ-vir(*terS*^S74F^) through the overexpression of either wild-type TerS (from a plasmid, pTerS) or cabRNA. Protein expression during infection prevented both cell death (Fig. 5A) as well as mutant phage propagation (Fig. 5B) through the generation of a structured cabRNA (Fig. 5C-D). Since the pTerS construct introduced into staphylococci cannot produce the activating RNA (it lacks the *terL*-encoded, downstream half of the cabRNA), these data suggest that the terminase small subunit protein is required for the generation an activating cabRNA with the proper length. Infection of staphylococci expressing a plasmid-encoded cabRNA also reduced the propagation of Φ80ɑ-vir(*terS*^S74F^) (Fig. S6G). Interestingly, in contrast to the plasmid-expressed cabRNA isolated from uninfected cells, a greater fraction of the RNA pulled down by the cyclase from staphylococci overexpressing the plasmid-borne cabRNA and infected with Φ80ɑ- vir(*terS*^S74F^) displayed an electrophoretic mobility consistent with the structured form of the cabRNA (Fig. S3D). This result supports our previous hypothesis that the secondary structures present in the inducer RNA, required for the activation of Ssc-CdnE03, are generated during phage infection.

## DISCUSSION

Here we show that the cyclic dinucleotidyltransferase of the *S. schleiferi* type I-B CBASS binds a 400-nucleotide RNA transcribed from the staphylococcal phages Φ80ɑ-vir and ΦNM1γ6 during infection, which we named cabRNA. This RNA forms secondary structures and interacts with a positively charged surface of the cyclase to promote the generation of cGAMP and initiate the CBASS anti-phage response. Most staphylococcal CBASS belong to the type I class and contain cyclases that are diverse phylogenetically, with many of them lacking the two positively charged residues at the first helix of the PβCD domain the surface exposed and conserved lysine or arginine residues along the spine and N-terminal domain that we identified as necessary for the interaction with the cabRNA (Fig. S7A). This suggests that there are other modes of cyclase activation, both in staphylococci and in other CBASS types, that do not sense viral RNA. This idea is also supported by our findings that the staphylococcal phages ΦNM4γ4 and Φ12γ3 do not produce cabRNA and are not restricted by Ssc-CBASS. Effective immunity against these phages may be provided by either the above-mentioned CBASS cyclases that sense viral signals different than the cabRNA, or by another, non-CBASS, mechanism of anti-phage defense. The positively charged surface made up of lysine and arginine residues in the N-terminal, PβCD, and C-terminal helices, however, are present in all members of the CdnE03 family of cyclases (Fig. S7B). These are widely distributed in different organisms and therefore we believe that the recognition of viral RNA for the activation of CBASS is a widespread mechanism across prokaryotes. Interestingly, the size of this surface (approximately 40 Å in length) should be able to only accommodate a dsRNA of approximately 20 base pairs, and therefore it remains incompletely understood which region of the cabRNA is most important for cyclase activation. We tested the two sequences with the strongest probability of hairpin formation within the cabRNA and found that only one mediated substantial, but not complete, activation (Fig. 3E). Therefore, it is possible that multiple Ssc-CdnE03 enzymes bind to the cabRNA to achieve high levels of cGAMP synthesis. This mechanism of activation is similar to that of human OAS1 and OAS3, which also require large dsRNA molecules for optimal activity ^39^. In the case of OAS1, the positively charged surface can interact with approximately with 18-20 base pairs (Fig. S4B). However, dsRNAs of this length can only provide limited activity (∼7-8% of the maximum) ^39^, therefore it is likely that longer species are also being recognized by OAS1 during viral infection *in vivo*. In OAS1, dsRNA binding leads to the rearrangement of the N and C lobes of the enzyme that positions the active site residues in the appropriate conformation for catalysis ^33^. Eukaryotic cyclases are activated by long, unmodified dsRNA in a sequence-independent manner, since since the presence of these molecules in the cytoplasm usually signals infection by RNA viruses ^15^. Given the lack of nuclear compartmentalization in bacteria, our results suggest that prokaryotic CBASS cyclases require a specific phage-derived structured RNA to avoid the autoimmunity that would be induced by host-derived transcripts.

It is not currently known how the cabRNA is produced during infection. Given that expression of a cabRNA in uninfected hosts results generates mostly the unfolded version of this species (Fig. S3D), which is pulled down by the Ssc-CdnE03 cyclase but fails to activate the CBASS response (Fig. S3C), we speculate that phage infection is required for the folding of the cabRNA into its cyclase-activating form. In addition, the S74F mutation in TerS, which does not change the cabRNA sequence, leads to the production of a longer, non-activating form of the cabRNA (Fig. 5D). This size change can be rescued *in trans* by the expression of wild-type TerS (Fig. 5D) and therefore we conclude that this protein is required to determine the proper length of the cabRNA. Interestingly, we and others ^35, 36^ found mutations in capsid proteins (Gp46 in the case of Φ80ɑ-vir) that enable escape from CBASS immunity. A structured cabRNA is produced during infection by these mutants (Fig. 5D) that is capable of fully activating the Ssc-CdnE03 cyclase (Fig S5E). In addition, compared to the *terS*^S74F^ mutant phages, the *gp46*^E105D^ escapers completely kill staphylococci in liquid cultures (Fig. S5C, compare to Fig. 5A) and form larger plaques in agar plates (Fig. S5A, compared to Fig. 5B). Therefore, we hypothesize that capsid mutations interfere with Ssc-CBASS immunity after cGAMP production. Finally, our pull-down assays also detected a host-derived RNA of the same length as the cabRNA. This RNA was not pulled down by Ssc-CdnE03 from total RNA of uninfected cells [nucleic acids were not detected after elution and agarose gel electrophoresis (Fig. 2B), nor by RT-PCR of the eluate (data not shown)]. Since it is not required for activation of cGAMP production *in vitro*, we speculate that the host RNA could perhaps be pulled down due to base-pair interactions with the cabRNA. However, given that these RNAs have identical lengths, we cannot rule out the possibility that the host RNA also has a function in Ssc-CBASS immunity *in vivo*. Future studies will elucidate the details of the biogenesis of the cabRNA and host-derived RNA during infection, as well as their molecular interactions with CdnE03 enzymes to initiate the synthesis of cyclic nucleotide second messengers.

## Acknowledgements

We would like to thank the members of the Marraffini laboratory for constructive feedback and encouragement, and Shelley Rankin at the University of Pennsylvania School of Veterinary Medicine for isolates of *S. schleiferi* 2142-05, 2317-03, and 5909-02. DVB is supported by an NIH Ruth L. Kirschstein NRSA F30 Individual Predoctoral Fellowship (F30AI157535) and an NIH Medical Scientist Training Program grant (T32GM007739) to the Weill Cornell/Rockefeller/Sloan Kettering Tri-Institutional MD-PhD Program. Support for this work comes from the National Institute of Health Director’s Pioneer Award 1DP1GM128184-01 to LAM. LAM is an investigator of the Howard Hughes Medical Institute.

## Authors contributions

Experiments were designed and analyzed by DVB, CR and LAM. CR and DVB conducted all experiments, except LC/MS of Ssc-CdnE03 products, which was performed by AMA. The paper was written by CR, DVB and LAM with the help and approval of all the authors.

## Competing interests

L.A.M. is a cofounder and Scientific Advisory Board member of Intellia Therapeutics, and a co-founder of Eligo Biosciences.

## Data and materials availability

all the data from these studies are available from the authors upon request.

## Supplementary Materials

### MATERIALS AND METHODS

#### Bacterial strains and growth conditions

The bacterial strains used in this study are listed in Supplementary Table S1. *Staphylococcus aureus* strain RN4220 ^23^ was grown at 37°C with shaking (220 RPM) in brain heart infusion (BHI) broth, supplemented with chloramphenicol (10 μg mL^-1^) or erythromycin (10 μg mL^-1^) to maintain pC194-based ^22^ or pE194-based plasmids ^40^, respectively. Cultures were supplemented with chloramphenicol (5 μg mL^-1^) to select for strains with chromosomally integrated Ssc-CBASS or Ssc-CdnE03. Gene expression was induced by the addition of 1 mM isopropyl-d-1-thiogalactopyranoside (IPTG) or 100 ng mL^-1^ anhydrotetracycline (aTc), where appropriate.

#### Bacteriophage propagation

The bacteriophages used in this study are listed in Supplementary Table S2. To generate a high-titer phage stock, an overnight culture of *S. aureus* RN4220 was diluted 1:100 and outgrown to mid-log phase (∼90 min) in BHI broth supplemented with 5 mM CaCl_2_. The culture was diluted to an optical density measurement at 600 nm (OD_600_) of 0.5 (∼1x10^8^ CFU mL^-1^). The culture was infected by adding phage at a multiplicity of infection (MOI) of 0.1 (∼1x10^7^ PFU mL^-1^), or by inoculating with either a single picked plaque or scrape of a frozen stock. The infected culture was grown at 37°C with shaking and monitored for lysis (full loss of turbidity was typically observed ∼3-4 hr). Culture lysates were centrifugated (4,300 x g for 10 min) to pellet cellular debris. The supernatant was collected, passed through a sterile membrane filter (0.45 μm), and stored at 4°C. Phage concentrations were determined by serially diluting the obtained stock in 10-fold increments and spotting 5 μL of each dilution on BHI soft agar mixed with RN4220 and supplemented with 5 mM CaCl_2_. After incubation overnight at 37°C, individual plaques (i.e. zones of no bacterial growth) were counted, and the viral titer was calculated.

#### Molecular cloning

The plasmids (and details of their construction) and the oligonucleotide primers used in this study are listed in Supplementary Tables S3 and S4, respectively. The coding sequences of Ssc-CBASS and phage gene products were obtained from genomic DNA preparations of *S. schleiferi* 2145-05 cultures ^20^ or phage stocks ^41^, respectively.

#### Chromosomal integration of Ssc-CBASS

Ssc-CBASS or Ssc-CdnE03, along with a chloramphenicol resistance (cmR) cassette, was integrated into the *hsdR* gene (which encodes the defective R-subunit of the restriction-modification system in *S. aureus* RN4220), an insertion site which was previously shown to not impact growth ^42^. Ssc-CBASS-cmR and Ssc-CdnE03-cmR were amplified from the plasmids pDVB303 and pDVB301 respectively, using primers oDVB565 and oDVB566, which were flanked with loxP sites at both ends followed by 60-bp homology regions to *hsdR*. Electrocompetent *S. aureus* RN4220 cells harboring the recombineering plasmid pPM300 were electroporated with 1-2 μg of PCR product and selected for with chloramphenicol (5 μg mL^-1^). Potential integrants were screened by colony PCR as well as for functional immunity, and then verified by Sanger sequencing.

#### Isolation of strictly lytic phage mutants

To construct a virulent mutant of the phage Φ80ɑ ^24^, we used a variation of a method previously described to generate ΦNM1γ6 ^25^, ΦNM4γ4 ^26^, and Φ12γ3 ^27^. Φ80ɑ-vir was isolated as a spontaneous escaper forming a clear plaque following Φ80ɑ infection of a BHI soft agar lawn of *S. aureus* RN4220 cells harboring plasmid pDVB08, which encodes a type III-A CRISPR-Cas system targeting the Φ80ɑ cI-like repressor. PCR of the Φ80ɑ-vir *cI* gene and Sanger sequencing confirmed an 8-bp deletion.

#### Soft agar phage infection

100 μL of an overnight bacterial culture was mixed with 5 mL BHI soft agar supplemented with 5 mM CaCl_2_ and poured onto BHI agar plates to solidify at room temperature (∼15 min). Phage lysates were serially diluted 10-fold and 4 μL was spotted onto the soft agar surface. Once dry, plates were incubated at 37°C overnight and visualized the next day. Individual plaques (zones of no bacterial growth) were enumerated manually.

#### Liquid culture phage infection

Overnight cultures were diluted 1:100 in BHI supplemented with 5 mM CaCl_2_ and the appropriate antibiotic for selection, outgrown at 37°C with shaking to mid-log phase (∼90 min), and normalized to OD_600_ 0.5. For the desired MOI, a calculated volume of phage stock was added to each culture and 150 μL was seeded into each well of a 96-well plate. OD_600_ was measured every 10 min in a microplate reader (TECAN Infinite 200 PRO) at 37°C with shaking.

#### RT-qPCR

Total RNA was extracted from *S. aureus* cells using a Direct-Zol^TM^ RNA MiniPrep Plus Kit (Cat. R2072). Extracted RNA was treated with TURBO^TM^ DNase (Thermo Fisher Scientific) before cDNA first-strand synthesis with SuperScript IV Reverse Transcriptase (Thermo Fisher Scientific) using random hexamers. qPCR was performed using Fast SYBR Green Master Mix (Life Technologies) and 7900HT Fast Real-Time PCR System (Applied Biosystems) with primer pairs for the *S. aureus* housekeeping gene *ptsG* (oDVB426/427), Ssc-CdnE03 (oDVB610/611), or Ssc-2TM (oDVB614/615).

#### Protein expression and purification

Ssc-CdnE03 and mutants were expressed and purified using the following approach: transformed BL21 (DE3) *E. coli* were grown in LB broth at 37°C with shaking to mid-log phase (OD_600_ 0.6-0.8), at which point the culture was cooled on ice for 10 min and induced with 0.2 mM IPTG for 16 hr at 18°C. Bacteria were harvested, resuspended in lysis buffer (25 mM Tris pH 7.4, 300 mM NaCl, 5% glycerol, 2 mM β-mercaptoethanol), and subjected to a single freeze-thaw cycle. The cells were incubated on ice with lysozyme, DNase I, and EDTA-free protease inhibitor cocktail. After incubating on ice for 40 min, the cells were lysed using sonication. Lysates were clarified by centrifugation and applied to cobalt affinity resin. After binding, the resin was washed extensively with lysis buffer prior to elution with lysis buffer containing 300 mM imidazole. Eluted proteins were then proteolyzed with TEV protease to remove the affinity tag during overnight 4°C dialysis to reaction buffer (25 mM HEPES-KOH pH 7.5, 250 mM KCl, 5% glycerol, 2 mM β-mercaptoethanol). The cleaved proteins were then passed over cobalt resin to collect the remaining tag (or uncleaved protein) and concentrated using 10,000 MWCO centrifugal filters (Amicon). Purified proteins were visualized by SDS-PAGE and used for downstream *in vitro* assays.

#### Nucleotide synthesis assays

Nucleotide synthesis assays were performed using a variation of the method described by Whiteley *et al.* ^9^. The final reactions (50 mM CAPSO pH 9.4, 50 mM KCl, 5 mM Mg(OAc)_2_, 1 mM DTT, 25 or 250 uM individual NTPs, trace amounts of [a-^32^P] NTP, 5 uM nucleic acid ligand, and 5 μM enzyme) were started with the addition of enzyme. All reactions except for those with RNA activator (2 hr) were incubated overnight at 37°C. For reactions with total RNA extracts, 500 ng was added to each condition. Reactions were stopped with the addition of 1 U of alkaline phosphatase, which removes triphosphates on the remaining NTPs and enables the visualization of cyclized nucleotide species. After a 1 hr incubation, 0.5 μL of the reaction was spotted 1.5 cm from the bottom of a PEI-cellulose thin-layer chromatography (TLC) plate, spaced 0.8 cm apart. TLC plates were developed in 1.5 M KH_2_PO_4_ pH 3.8 until the buffer front reached 1 cm from the top (∼12 cm). The TLC plates were completely dried, covered with plastic wrap and exposed to a phosphor screen before detection by a Typhoon Trio Imager System.

To purify the Ssc-CdnE03 cyclic nucleotide product for mass spectrometry analysis, nucleotide synthesis reaction conditions were scaled up to 1 mL reactions containing 5 uM Ssc-CdnE03, 250 uM ATP, 250 uM GTP, approximately 5 ng of cabRNA, in 50 mM CAPSO pH 9.4, 50 mM KCl, 5 mM Mg(OAc)2, 1 mM DTT buffer. Reactions were incubated with gentle shaking for 24 hr at 37°C followed by Quick CIP (NEB) treatment for 4 hr at 37°C. Following incubation, reactions were filtered through a 10,000 MWCO centrifugal filter (Amicon) to remove protein.

#### Nucleotide High Resolution Mass Spectrometry Analysis

All solvents and reagents used for chromatography were LC-MS grade. UPLC-HRMS data was acquired on a Sciex ExcionLC UPLC coupled to an X500R mass spectrometer, controlled by SCIEXOS software. Chromatography was carried out on a Phenomenex Acquity UPLC® BEH Shield RP18 (2.1 x 150 mm, 1.7 µm), under the following conditions: 3% B from 0.0 to 6.0 min, from 3% to 10% B from 6.0 to 16.0 min, 10% B until 18.0, 95% B from 18.0 to 21.0 and 3% B from 22.0 to 27.0 (A: water + 0.1% formic acid; B: acetonitrile + 0.1% formic acid) (pending of new buffer), with a flow rate of 0.25 mL/min and 1µL of injection volume. HRMS analysis were performed in positive and negative electrospray ionization mode in the range m/z 100-1200 for MS1 and MS2 scans; the maximum candidate ions subjected for Q2-MS2 experiments was 7, declustering potential of 80 V, collision energy of 5 V and temperature of 500°C. For ESI+ HRMS experiments the spray voltage was set in 5500 V, the Q2 collision energy at 30 V with a spread of 10 V, whereas the spray voltage for ESI-HRMS was set in 4500V and the Q2 collision energy at 35 V with a spread of 10 V. The concentration of the standard solutions was 6.25 µM, and all the solutions were centrifuged (13,000 rpm x 3 min) before injection. The molecular ions for ESI modes were analyzed for all compounds, but the fragmentation in ESI-mode showed a better consistency and was consequently used for the structural analysis. The data analysis was carried out with MestReNova software (14.3.0), data output was converted with MSConvert from Proteowizard, MS2 mirror plot was obtained from GNPS using averaged MS2 spectra from GNPS molecular networking.

#### RNA extraction from phage infection

10 mL of a mid-log phase *S. aureus* RN4220 culture normalized to OD_600_ 0.5 was infected with phage at MOI 10. Infection was allowed to proceed for 30 min, just before the completion of the first burst. Cells were pelleted at 4,300 x g for 5 min and flash-frozen with liquid nitrogen. The pellet was resuspended in 150 μL PBS and 50 μL lysostaphin (5 mg mL^-1^) and incubated at 37°C for 30 min. Total RNA was extracted from *S. aureus* cells using a Direct-Zol^TM^ RNA MiniPrep Plus Kit (Cat. R2072). Briefly, 450 μL Trizol was added to the lysate, vigorously vortexed and centrifuged at 16,000 x g for 30 seconds. 650 μL of 100% ethanol was added to the supernatant and the samples were thoroughly vortexed. The entire volume was passed through a Zymo-Spin IIICG Column followed by in-column treatment with DNase I for 15 min at room temperature. The column was washed according to the manufacturer’s protocol and RNA was eluted in 100 μL nuclease-free water.

#### RNA pull-down assay

His6-MBP-tagged Ssc-CdnE03 was expressed and purified as described above. Purified His6-MBP tag alone was prepared alongside as a negative control. After immobilizing ∼0.2 mg of protein on cobalt resin, the column was washed extensively with lysis buffer prior to the addition of 5 mL of lysis buffer containing 1 mM MgCl_2_, 5 units of RNaseOUT^TM^ (ThermoFisher, Cat. 10777019), and 100 μg of total RNA extracted from cultures with or without phage infection. The RNA was incubated with the tagged Ssc-CdnE03 on the column for 40 min before washing the column with 5 volumes of lysis buffer. The column was treated with His6-tagged TEV protease to release the Ssc-CdnE03 and bound RNA. Eluted protein was collected for each sample and combined with TRI Reagent (Zymo Research, Cat. R2050-1-200). RNA was then extracted according to the Direct-Zol^TM^ RNA MiniPrep Plus Kit (Cat. R2072) manufacturer’s protocol. The final RNA product was run on a 2% agarose 1X TAE gel and stained with SyBr Gold or ethidium bromide. Eluted protein samples were collected as controls for visualization by SDS-PAGE.

#### RNA-sequencing

Reverse transcription of the RNA isolated from the pull-down assay was performed as detailed above. Briefly, RNA was treated with TURBO^TM^ DNase (Thermo Fisher Scientific) before cDNA first-strand synthesis with SuperScript IV Reverse Transcriptase using random hexamers. Second-strand synthesis of the cDNA was performed with Q5 DNA polymerase at 15°C for 2 hr, followed by 75°C for 10 min in the presence of RNase H and DMSO. The cDNA was then sheared to 150-bp fragments using an S220 Covaris Focused-Ultrasonicator (peak incident power: 175 W, duty factor: 10%, cycles per burst: 200, treatment time: 430 s, temperature 4°C) in S-Series Holder microTUBEs (PN 500114). Library preparation was performed using the Illumina TruSeq Library Prep Kit. Quantification and quality check of DNA libraries were confirmed by Qubit 4.0 Fluorometer and Agilent Bioanalyzer/Tapestation, respectively. loaded to a flow cell on an Illumina MiSeq instrument (paired ends, 150 cycles). 12 pM DNA library was loaded on an Illumina MiSeq instrument for paired-end sequencing (2 x 150 cycle). Bowtie2 via the Galaxy open-source interface ^43^ was used to align sequencing reads to phage and host genomes. A custom Python script was used to convert the output SAM alignments into CSV files.

#### RNA structure prediction

RNA secondary structures were analyzed using the ViennaRNA 2.0 package ^30^ and visualized via the SnapGene interface.

#### *In vitro* transcription of cabRNA

*In vitro* transcription (IVT) was performed according to the Thermo Scientific TranscriptAid T7 High Yield Transcription Kit protocol (Cat. K0441). Linear dsDNA for the cabRNA, host pull-down RNA, and *terS^S74F^* phage escaper RNA sequences were PCR-amplified using oCR190/193 (sense cabRNA), oCR191/192 (antisense cabRNA), oCR194/197 (sense host RNA), oCR195/196 (antisense host RNA), and oDVB691/oCR193 (*terS^S74F^*phage escaper RNA). The target sequence was placed downstream of a T7 promoter, which was inverted for antisense transcription reactions. For high yield *in vitro* transcription reactions, 1 μg of PCR product was combined with TranscriptAid Enzyme mix and NTPs. Following a 4 hr incubation period at 37°C, transcripts were purified according to the Direct-Zol^TM^ RNA MiniPrep Plus Kit (Cat. R2072) manufacturer’s protocol. To stimulate the re-folding and formation of a structured RNA product, the purified IVT samples were heated at 95°C for 5 min in a heat block, which was slowly cooled down to room temperature over 1 hr. Where indicated, IVT products were either heat-treated (“folded”) or untreated.

#### Structural prediction and analysis of Ssc-CdnE03

The amino acid sequence of Ssc-CdnE03 sequence was used to seed a position-specific iterative BLAST (PSI-BLAST) search of the NCBI non-redundant protein and conserved domain databases (composition-based adjustment, E-value threshold 0.01). Putative domains identified from this search include a C-terminal “nucleotidyltransferase (NT) domain of 2’5’-oligoadenylate (2-5A) synthetase” (NT_2-5OAS) domain (Residues 61-204; E-value 2.63e-15) and an N-terminal “tRNA nucleotidyltransferase” (CCA-adding enzyme) domain (Residues 5-158; E-value 8.01e-04). A structure of the Ssc-CdnE03 was predicted using AlphaFold (ColabFold). Following structure determination, pairwise structural comparison of the rank 1 model to the full PDB database was performed using DALI. The ConSurf database was used to visualize conserved structural features of the Ssc-CdnE03. Structural alignments and generation of surface electrostatics with apo-OAS1 (PDB:4RWQ) and OAS1:dsRNA (PDB:4RWO) were performed using PyMOL.

#### Generation and isolation of escaper bacteriophages

Overnight cultures of *S. aureus* RN4220 were diluted 1:100 and outgrown at 37°C with shaking for 1 hr, infected with Φ80ɑ-vir (MOI 1) for 20 min, and then treated with 1% ethyl methanesulfonate (EMS), a chemical mutagen. Cultures were allowed to lyse for 3 hr before pelleting debris and sterile-filtering the supernatant to obtain an EMS-treated mutant phage library. 100 μL of RN4220 overnight cultures harboring Ssc-CBASS were infected with a high titer mutant phage library in BHI soft agar and then plated. After incubating at 37°C overnight, individual phage plaques were picked from the top agar and resuspended in 50 μL of BHI liquid medium. Phage lysates were further purified over two rounds of passaging on RN4220 harboring Ssc-CBASS.

#### Whole-genome sequencing and analysis

Genomic DNA from high-titer phage stocks was extracted using a previously described method ^41^. DNA was sheared to 300-bp fragments using an S220 Covaris Focused-Ultrasonicator (peak incident power: 140 W, duty factor: 10%, cycles per burst: 200, treatment time: 80 s, temperature 4°C) in S-Series Holder microTUBEs (PN 500114). Library preparation was performed using the Illumina Nextera XT DNA Library Preparation Kit protocol (Cat. FC-131-1096). 12 pM of the library was loaded on an Illumina MiSeq instrument for paired-end sequencing (2 x 150 cycle). Bowtie2 via the Galaxy open-source interface ^43^ was used to align sequencing reads to phage and host genomes. A custom Python script was used to convert the output SAM alignments into CSV files.

#### Generation of recombinant ΦNM1γ6 *terS^S74F^* mutants

Wild-type ΦNM1γ6 was passaged on *S. aureus* RN4220 harboring Ssc-CBASS and pTerS or pTerS^S74F^ to enable recombination. The infected culture supernatant was spotted onto a lawn of RN4220 with *Ssc*CBASS in BHI soft agar to isolate individual escaper plaques. The *terS* gene was amplified by PCR and the S74F mutation was confirmed by Sanger sequencing.

#### Phylogenetic analysis of CD-NTase sequences

The CD-NTases from *S. schleiferi* are most similar to the CdnE subtype 03 (CdnE03) described by Whiteley *et al.* ^9^. All CD-NTase enzymes were aligned using TCoffee Multiple Sequence Alignment tool (default parameters) and used to construct a phylogenetic tree with Geneious Prime using the neighbor-joining method and Jukes-Cantor genetic distance model with no outgroup.

### SUPPLEMENTARY SEQUENCES FILE

#### Sequences of RNA species pulled down from Ssc-CdnE03-RNA pull-down assay (related to Figures 2D and S2G-H)

**400-bp bacteriophage RNA during <λ>80α-vir infection (cabRNA_<λ>80α_):** 5’-

AGAGGAGAACCUCAAGAGGCUUACAGUAAGAAAUAUGACCAUUUAAACGAUGAA GUGGAAAAAGAGGUUACUUACACAAUCACACCAACUUUUGAAGAGCGUCAGAGA UCUAUUGACCACAUACUAAAAGUUCAUGGUGCGUAUAUCGACAAAAAAGAAAUUA CUCAGAAGAAUAUUGAGAUUAAUAUUGGUGAGUACGAUGACGAAAGUUAAAUUA AACUUUAACAAACCAUCUAAUGUUUUCAACAGAAACAUAUUCGAAAUACUAACCA AUUACGAUAACUUCACUGAAGUACAUUACGGUGGAGGUUCGAGUGGUAAGUCUC ACGGCGUUAUACAAAAAGUUGUACUUAAAGCAUUGCAAGACUGGAAAUAUCCUA GGCGUAUACUAUGGCUUAGA-3’

**1237-bp bacteriophage RNA during <λ>80α-vir *gp40*_S74F_ infection:** 5’-

AUGACUAAAAAGAAAUAUGGAUUAAAAUUAUCAACAGUUCGAAAGUUAGAAGAUG AGUUGUGUGAUUAUCCUAAUUAUCAUAAGCAACUCGAAGAUUUAAGAAGUGAAA UAAUGACACCAUGGAUUCCAACAGAUACAAAUAUAGGCGGGGAGUUUGUACCGU CUAAUACAUCGAAAACAGAAAUGGCAGUAACUAAUUAUCUUUGUAGUAUACGAAG AGGUAAAAUCCUUGAGUUUAAGAGCGCUAUUGAACGUAUAAUCAACACAUCAAG UAGGAAAGAACGCGAAUUCAUUCAAGAGUAUUAUUUUAAUAAAAAGGAAUUAGU GAAAGUUUGUGAUGACAUACACAUUUCUGAUAGAACUGCUCAUAGAAUCAAAAG GAAAAUCAUAUCUAGAUUGGCGGAAGAGUUAGGGGAAGAGUGAAAUUGGCAGUA AAGUGGCAGUUUUUGAUACCUAAAAUGAGAUAUUAUGAUAGUGUAGGAUAUUGA CUAUCUUACUGCGUUUCCCUUAUCGCAAUUAGGAAUAAAGGAUCUAUGUGGGUU GGCUGAUUAUAGCCAAUCCUUUUUUAAUUUUAAAAAGCGUAUAGCGCGAGAGUU GGUGGUAAAUGAAAUGAACGAAAAACAAAAGAGAUUCGCAGAUGAAUAUAUAAUG AAUGGAUGUAAUGGUAAAAAAGCAGCAAUUUCAGCAGGUUAUAGUAAGAAAACA GCAGAGUCUUUAGCAAGUCGAUUGUUAAGAAAUGUUAAUGUUUCGGAAUAUAUU AAAGAACGAUUAGAACAGAUACAAGAAGAGCGUUUAAUGAGCAUUACAGAAGCU UUAGCGUUAUCUGCUU**U**UAUUGCUAGAGGAGAACCUCAAGAGGCUUACAGUAAG AAAUAUGACCAUUUAAACGAUGAAGUGGAAAAAGAGGUUACUUACACAAUCACAC CAACUUUUGAAGAGCGUCAGAGAUCUAUUGACCACAUACUAAAAGUUCAUGGUG CGUAUAUCGACAAAAAAGAAAUUACUCAGAAGAAUAUUGAGAUUAAUAUUGGUGA GUACGAUGACGAAAGUUAAAUUAAACUUUAACAAACCAUCUAAUGUUUUCAACAG AAACAUAUUCGAAAUACUAACCAAUUACGAUAACUUCACUGAAGUACAUUACGGU GGAGGUUCGAGUGGUAAGUCUCACGGCGUUAUACAAAAAGUUGUACUUAAAGCA UUGCAAGACUGGAAAUAUCCUAGGCGUAUACUAUGGCUUAGA-3’

400-bp **bacteriophage RNA during <λ>NM1ψ6 infection (cabRNA_<λ>NM1ψ6_):** 5’-

AGAGGAGAACCUCAAGAGGCUUACAGUAAGAAAUAUGACCAUUUAAACGAUGAA GUGGAAAAAGAGGUUACUUACACAAUCACACCAACUUUUGAAGAGCGUCAGAGA UCUAUUGACCACAUACUAAAAGUACAUGGUGCGUAUAUCGAUAAAAAAGAAAUUA CUCAGAAGAAUAUUGAGAUUAAUAUUGGUGAGUACGAUGACGAAAGUUAAAUUA AACUUUAACAAACCAUCUAAUGUUUUCAAUAGAAACAUAUUCGAAAUACUAACCA AUUACGAUAACUUCACUGAAGUACAUUACGGUGGAGGUUCGAGCGGUAAGUCUC ACGGCGUUAUACAAAAAGUUGUACUUAAAGCAUUGCAAGACUGGAAAUAUCCUA GGCGUAUACUAUGGCUUAGA-3’

**400-bp host RNA during<Ι>80α-vir infection:** 5’-

ACAUUAACGACAACUCAAGGUAUUCCAAUUAAUAUUAGAGGGCAAAUUGACCGUAUCGAUACGUAUACAAAGAAUGAUACAAGUUUUGUUAAUAUCAUUGACUAUAAAUCCUCUGAAGGUAGUGCGACACUUGAUUUAACGAAAGUAUAUUAUGGUAUGCAAAUGCAAAUGAUGACAUACAUGGAUAUCGUUUUACAAAAUAAACAACGCCUUGGAUUAACAGAUAUUGUGAAACCAGGUGGAUUAUUAUACUUCCAUGUACAUGAACCUAGAAUUAAAUUUAAAUCAUGGUCUGAUAUUGAUGAAGAUAAACUAGAACAAGAUUUAAUUAAAAAGUUUAAGUUGAGUGGUUUAGUUAAUGCAGACCAAACUGUUAUUGAUGCAUUGGAUAUUCGUUUAG-3’

**1237-bp host RNA during <Ι>80α-vir (*gp40*_S74F_) infection:** 5’-

CAGAUGGAUGAAGCAUUUGUUUGUUAUGUUGCUAUGACUAGAGCUAAGGGAGAUGUUACAUUUUCUUACAGUCUAAUGGGAUCAAGUGGUGAUGAUAAGGAGAUCAGCCCAUUUUUAAAUCAAAUUCAAUCAUUGUUCAACCAAUUGGAAAUUACUAACAUUCCUCAAUACCAUGAAGUUAACCCAUUGUCACUAAUGCAACAUGCUAAGCAAACCAAAAUUACAUUAUUUGAAGCAUUGCGUGCUUGGUUAUAUGAUGAAAUUGUGGCUGAUAGUUGGUUAGAUGCUUAUCAAGUAAUUAGAGAUAGCGAUCAUUUAAAUCAAGGUUUAGAUUAUUUAAUGUCAGCAUUAACGUUUGACAAUGAAACUGUAAAAUUAGGUGAAACGUUGUCUAAAGAUUUAUAUGGUAAGGAAAUCAAUGCCAGUGUAUCCCGUUUUGAAGGUUAUCAACAAUGCCCAUUUAAACACUAUGCGUCACAUGGUCUGAAACUAAAUGAGCGAACGAAGUAUGAACUUCAAAACUUUGAUUUAGGUGAUAUUUUCCAUUCUGUUUUAAAAUAUAUAUCUGAACGUAUUAAUGGCGAUUUUAAACAAUUAGACCUGAAAAAAAUAAGACAAUUAACGAAUGAAGCAUUGGAAGAAAUUUUACCUAAAGUUCAGUUUAAUUUAUUAAAUUCUUCAGCUUACUAUCGUUAUUUAUCAAGACGCAUUGGCGCUAUUGUAGAAACAACACUAAGCGCAUUAAAAUAUCAAGGCACGUAUUCAAAGUUUAUGCCAAAACAUUUUGAGACAAGUUUUAGAAGGAAACCAAGAACAAAUGACGAAUUAAUUGCACAAACAUUAACGACAACUCAAGGUAUUCCAAUUAAUAUUAGAGGGCAAAUUGACCGUAUCGAUACGUAUACAAAGAAUGAUACAAGUUUUGUUAAUAUCAUUGACUAUAAAUCCUCUGAAGGUAGUGCGACACUUGAUUUAACGAAAGUAUAUUAUGGUAUGCAAAUGCAAAUGAUGACAUACAUGGAUAUCGUUUUACAAAAUAAACAACGCCUUGGAUUAACAGAUAUUGUGAAACCAGGUGGAUUAUUAUACUUCCAUGUACAUGAACCUAGAAUUAAAUUUAAAUCAUGGUCUGAUAUUGAUGAAGAUAAACUAGAACAAGAUUUAAUUAAAAAGUUUAAGUUGAGUGGUUUAGUUAAUGCAGACCAAACUGUUAUUGAUGCAUUGGAUAUUCGUUUAG-3’

#### Sequences of *gp40* from bacteriophages used in this study

***gp40* sequence from <Ι>80α-vir:**

5’-

ATGAACGAAAAACAAAAGAGATTCGCAGATGAATATATAATGAATGGATGTAATGGT AAAAAAGCAGCAATTTCAGCAGGTTATAGTAAGAAAACAGCAGAGTCTTTAGCAAG TCGATTGTTAAGAAATGTTAATGTTTCGGAATATATTAAAGAACGATTAGAACAGAT ACAAGAAGAGCGTTTAATGAGCATTACAGAAGCTTTAGCGTTATCTGCTTCTATTGC TAGAGGAGAACCTCAAGAGGCTTACAGTAAGAAATATGACCATTTAAACGATGAAG TGGAAAAAGAGGTTACTTACACAATCACACCAACTTTTGAAGAGCGTCAGAGATCT ATTGACCACATACTAAAAGTTCATGGTGCGTATATCGACAAAAAAGAAATTACTCAG AAGAATATTGAGATTAATATTGGTGAGTACGATGACGAAAGTTAA-3’

***gp40* sequence from <Ι>80α-vir *gp40*_S74F_:** 5’-

ATGAACGAAAAACAAAAGAGATTCGCAGATGAATATATAATGAATGGATGTAATGGT AAAAAAGCAGCAATTTCAGCAGGTTATAGTAAGAAAACAGCAGAGTCTTTAGCAAG TCGATTGTTAAGAAATGTTAATGTTTCGGAATATATTAAAGAACGATTAGAACAGAT ACAAGAAGAGCGTTTAATGAGCATTACAGAAGCTTTAGCGTTATCTGCTT**T**TATTGC TAGAGGAGAACCTCAAGAGGCTTACAGTAAGAAATATGACCATTTAAACGATGAAG TGGAAAAAGAGGTTACTTACACAATCACACCAACTTTTGAAGAGCGTCAGAGATCT ATTGACCACATACTAAAAGTTCATGGTGCGTATATCGACAAAAAAGAAATTACTCAG AAGAATATTGAGATTAATATTGGTGAGTACGATGACGAAAGTTAA-3’

***gp40* sequence from <Ι>NM1ψ6:** 5’-

ATGAACGAAAAACAAAAGAGATTCGCAGATGAATATATAATGAATGGATGTAATGGT AAAAAAGCAGCAATTACAGCAGGTTATAGTAAGAAAACAGCAGAGTCTTTAGCAAG TCGATTGTTAAGAAATGTTAATGTTTCGGAATATATTAAAGAACGATTAGAACAGAT ACAAGAAGAGCGTTTAATGAGTATTACAGAAGCTTTAGCGTTATCTGCTTCTATTGC TAGAGGAGAACCTCAAGAGGCTTACAGTAAGAAATATGACCATTTAAACGATGAAG TGGAAAAAGAGGTTACTTACACAATCACACCAACTTTTGAAGAGCGTCAGAGATCT ATTGACCACATACTAAAAGTACATGGTGCGTATATCGATAAAAAAGAAATTACTCAG AAGAATATTGAGATTAATATTGGTGAGTACGATGACGAAAGTTAA-3’

***gp40* sequence from <Ι>NM1ψ6 *gp40*_S74F_ recombinant:** 5’-

ATGAACGAAAAACAAAAGAGATTCGCAGATGAATATATAATGAATGGATGTAATGGT AAAAAAGCAGCAATTTCAGCAGGTTATAGTAAGAAAACAGCAGAGTCTTTAGCAAG TCGATTGTTAAGAAATGTTAATGTTTCGGAATATATTAAAGAACGATTAGAACAGAT ACAAGAAGAGCGTTTAATGAGCATTACAGAAGCTTTAGCGTTATCTGCTT**T**TATTGC TAGAGGAGAACCTCAAGAGGCTTACAGTAAGAAATATGACCATTTAAACGATGAAG TGGAAAAAGAGGTTACTTACACAATCACACCAACTTTTGAAGAGCGTCAGAGATCT ATTGACCACATACTAAAAGTTCATGGTGCGTATATCGACAAAAAAGAAATTACTCAG AAGAATATTGAGATTAATATTGGTGAGTACGATGACGAAAGTTAA-3’

### SUPPLEMENTARY TABLES

**Table S1.** Bacterial strains used in this study.

| **Species** | **Strain** | **Genotype** | **Origin** |
| --- | --- | --- | --- |
| *S. aureus* | RN4220 | Wild type | Kreiswerth et al., Nature (1983) |
| *S. aureus* | RN4220 | ::Ssc-CdnE03-cmR | Chromosomal integration (see Methods) |
| *S. aureus* | RN4220 | ::Ssc-CBASS-cmR | Chromosomal integration (see Methods) |
| *E. coli* | BL21 (DE3) | F<sup>-</sup> *ompT gal dcm lon hsdS<sub>B</sub> (r<sub>B</sub><sup>-</sup>m<sub>B</sub><sup>-</sup>)* | Thermo Fisher Scientific |

**Table S2.** Phages used in this study.

| Phage | Host | Genotype | Origin |
| --- | --- | --- | --- |
| Φ80α-vir | *S. aureus* | Wild type | This study; strictly lytic mutant of Φ80α isolated from type III CRISPR-Cas targeting of *cI* repressor gene |
| Φ80α-vir | *S. aureus* | *terS* S74F (C221>T) | This study; isolated from EMS chemical mutagenesis screen for Ssc-CBASS escapers |
| Φ80α-vir | *S. aureus* | *gp46* E105D (G315>T) | This study; isolated from EMS chemical mutagenesis screen for Ssc-CBASS escapers |
| Φ80α-vir | *S. aureus* | *gp46* E105K (G313>A) | This study; isolated from EMS chemical mutagenesis screen for Ssc-CBASS escapers |
| Φ80α-vir | *S. aureus* | *gp46* R110H (G329>A) | This study; isolated from EMS chemical mutagenesis screen for Ssc-CBASS escapers |
| ΦNM1γ6 | *S. aureus* | Wild type | Goldberg et al., Nature (2014) |
| ΦNM1γ6 | *S. aureus* | Φ80α *terS* S74F | This study; isolated through recombination with Φ80α-vir *terS*<sup>S74F</sup> |
| ΦNM4γ4 | *S. aureus* | Wild type | Heler et al., Nature (2015) |
| Φ12γ3 | *S. aureus* | Wild type | Modell et al., Nature (2017) |

**Table S3.**
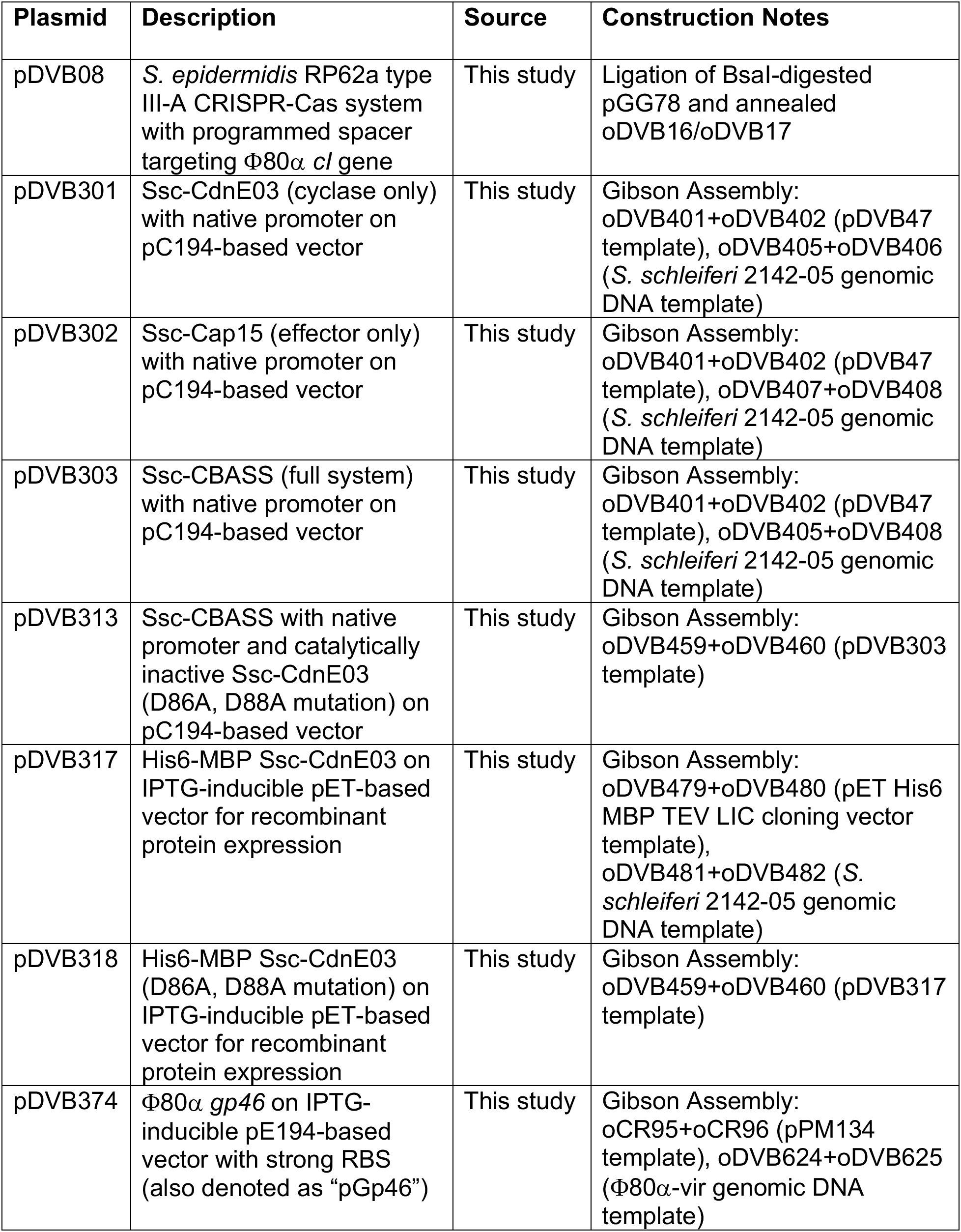
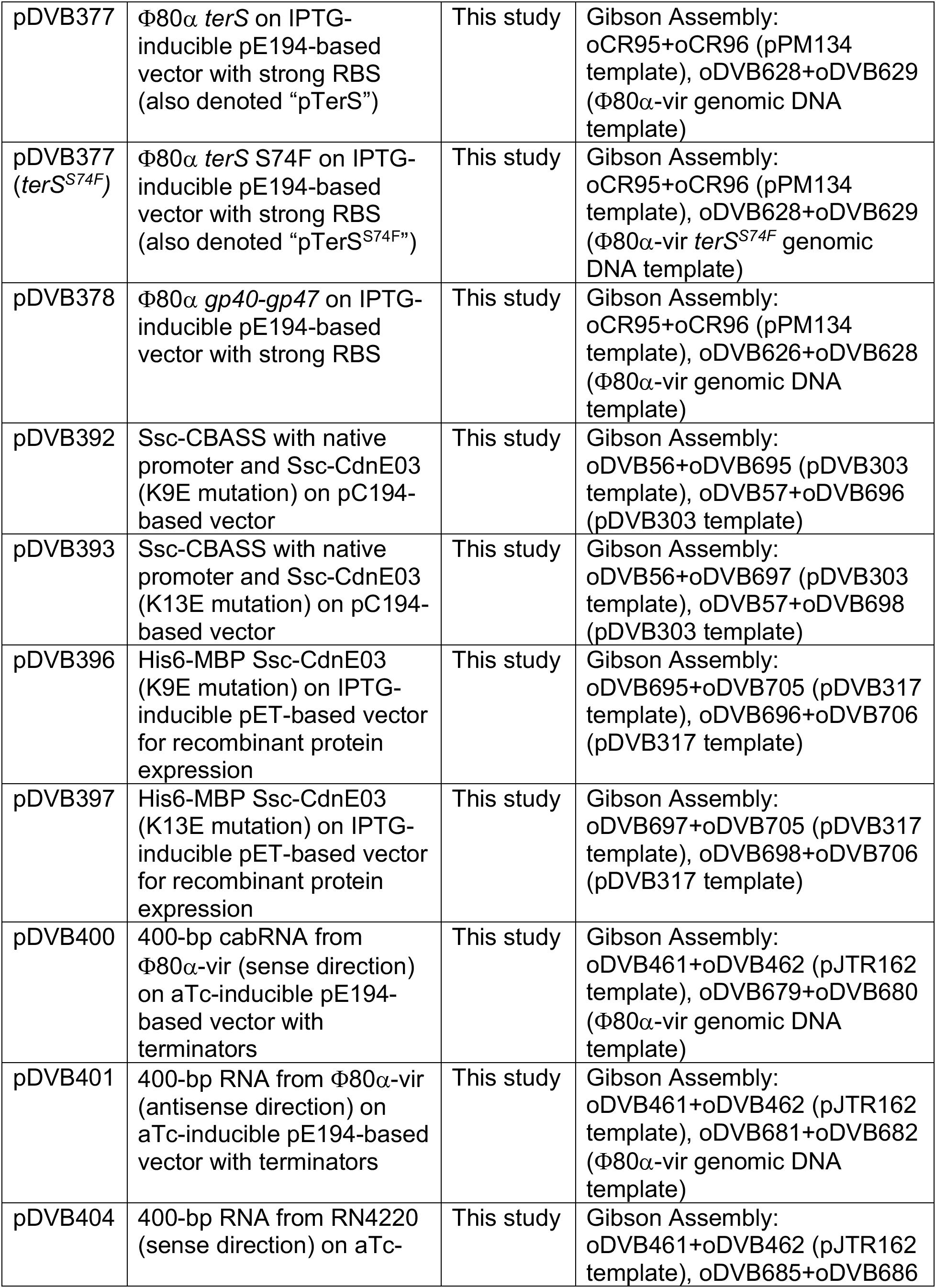
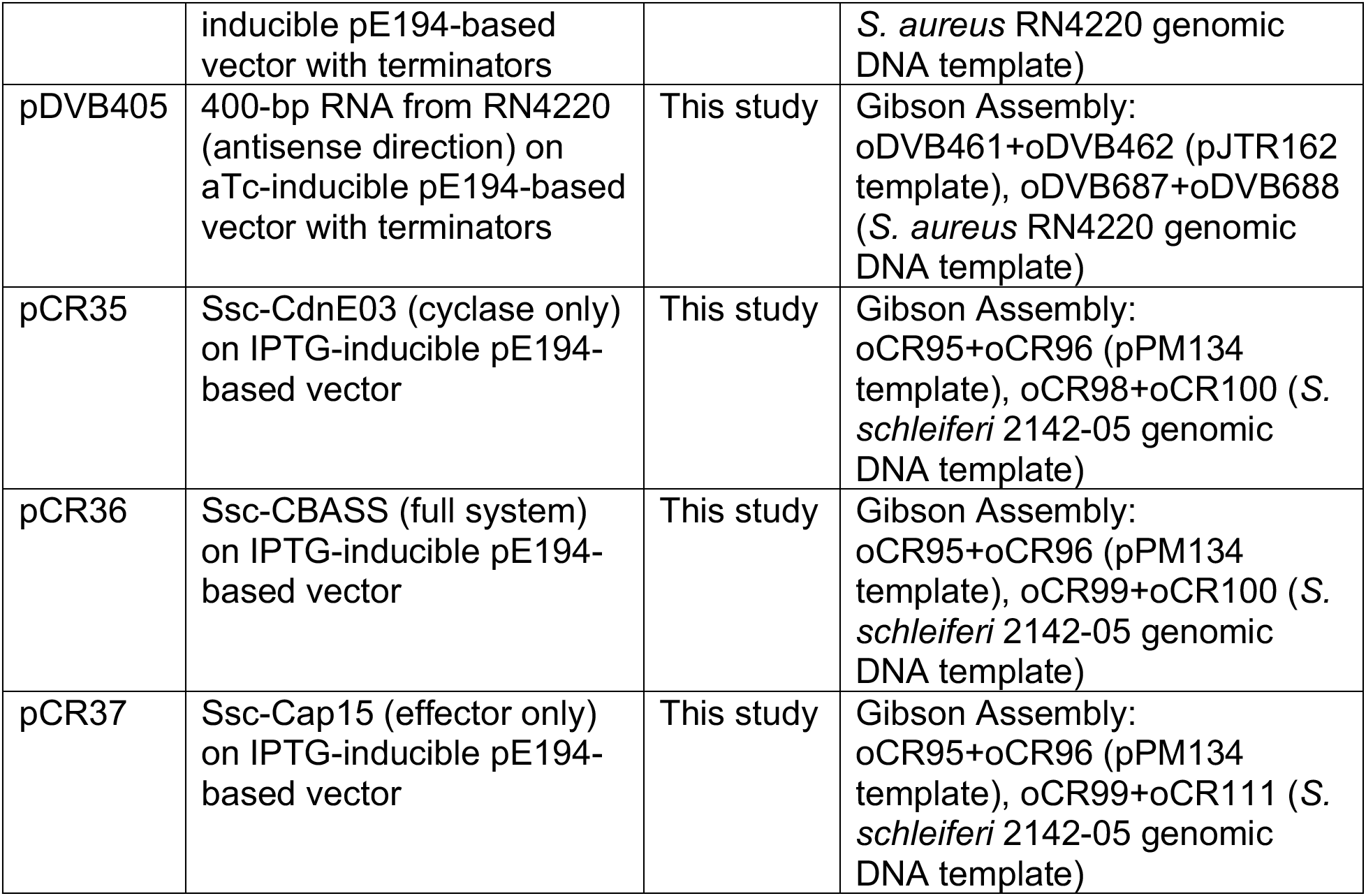
Plasmids used in this study.

**Table S4.**
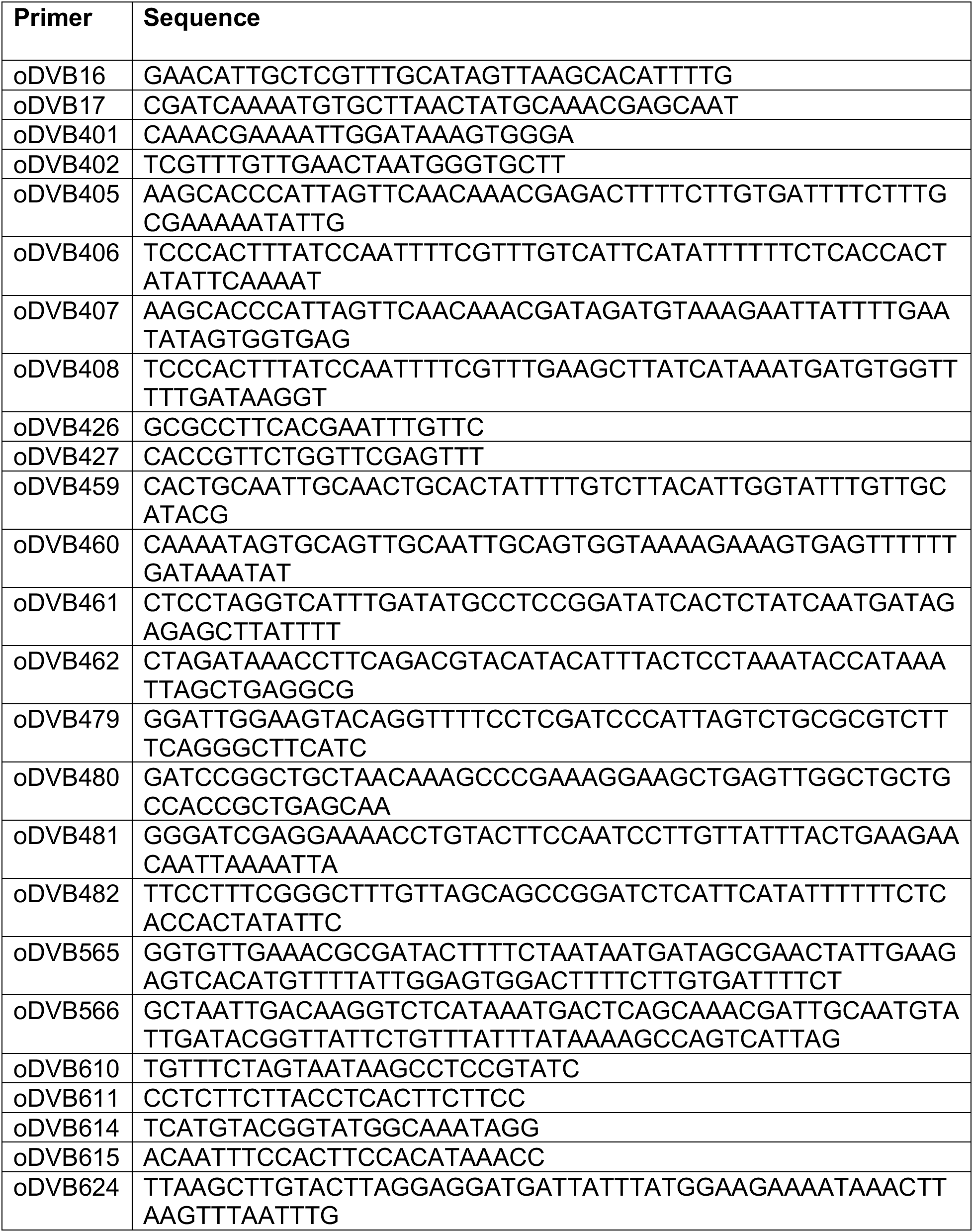
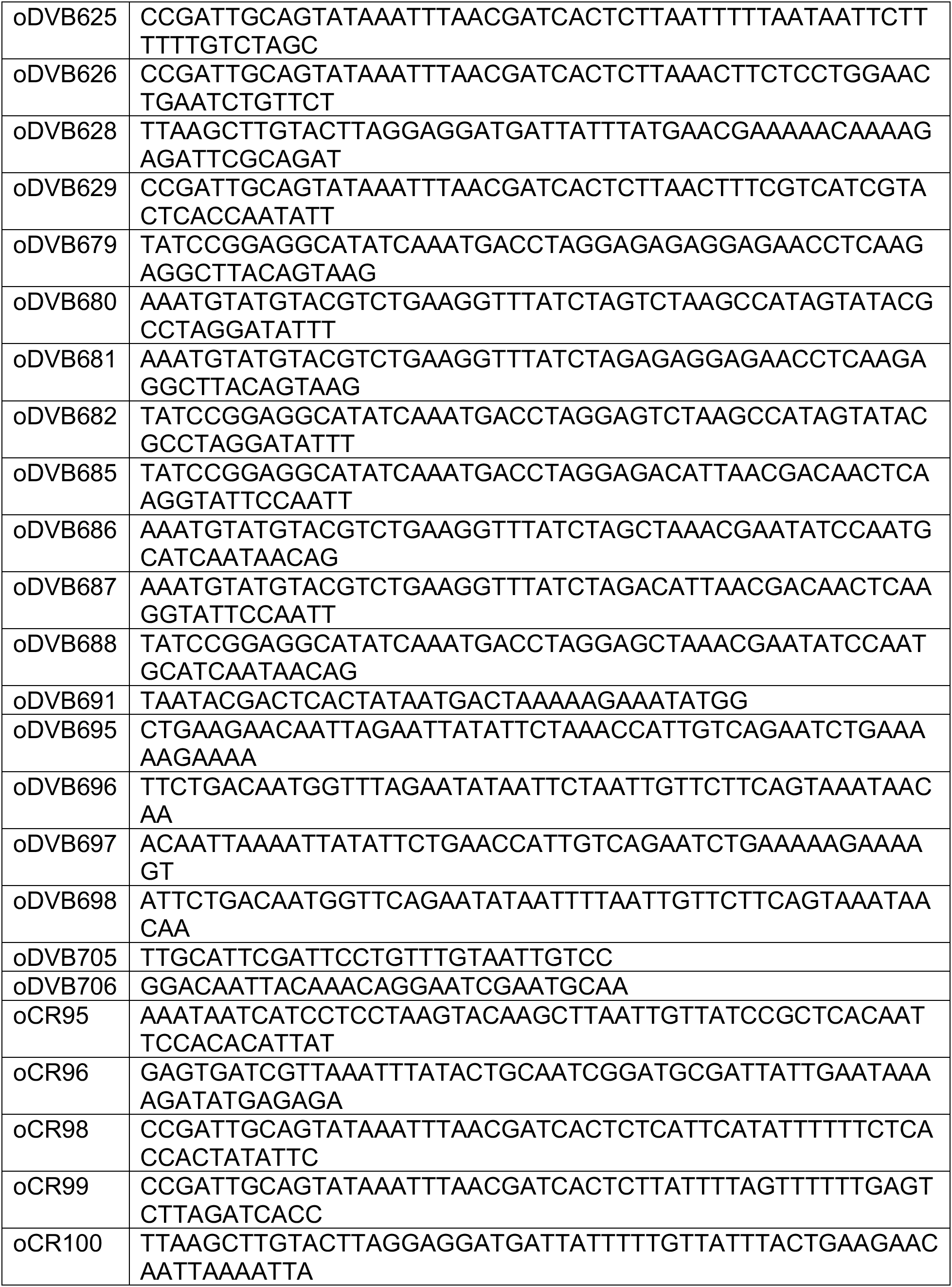
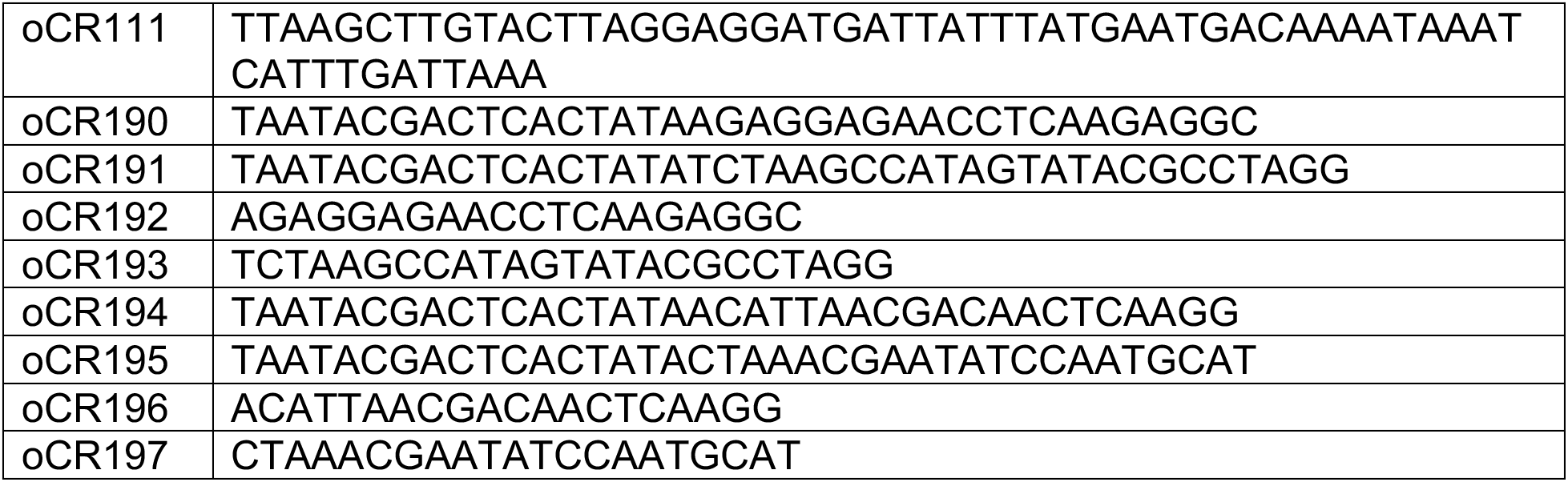
Oligonucleotide primers used in this study.

**Table S5.**
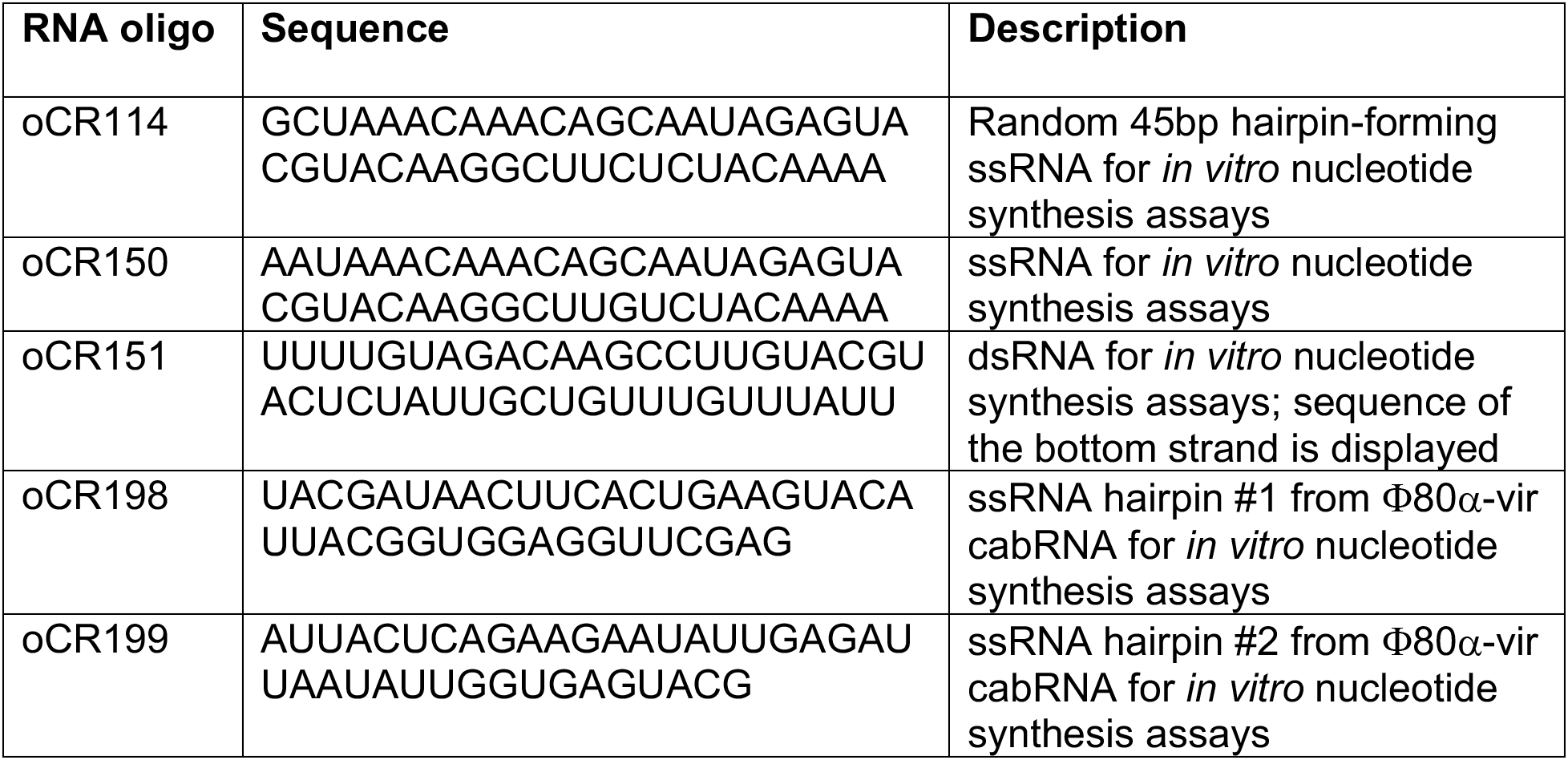
RNA oligonucleotides used in this study.

### Supplementary Text

#### High Resolution Mass Spectrometry Analysis of Ssc-CdnE03 reaction products

The Full HRMS of the products of the Ssc-CdnE03 cyclase shows that this peak corresponds to a compound identified from its mono and double charged ions in the positive and negative ionization modes, that allow the molecular formula C_15_H_19_N_5_O_13_P_2_ to be predicted, consistent with the molecular formula of cGAMP.

MS^2^ experiments were run in both ionization modes, however, the negative shows more diagnostic fragments and a consistent fragmentation pattern throughout the set of evaluated ions. The MS^2^ spectra of both cGAMP isomers show identical fragmentation to the Ssc-CdnE03 product, being particularly relevant the presence of fragment ions from cleavages a (a1, a2), b (b1, b2) and c (c1, c2) that allow identifying this product as a dimeric nucleotide constituted by GMP and AMP units. Based on these results we conclude that the Ssc-CdnE03 product is a cGAMP isomer or a mixture of them.

##### NEGATIVE ION MODE ACQUISITION

**Characterization of standard 3’,3’-cGAMP:**

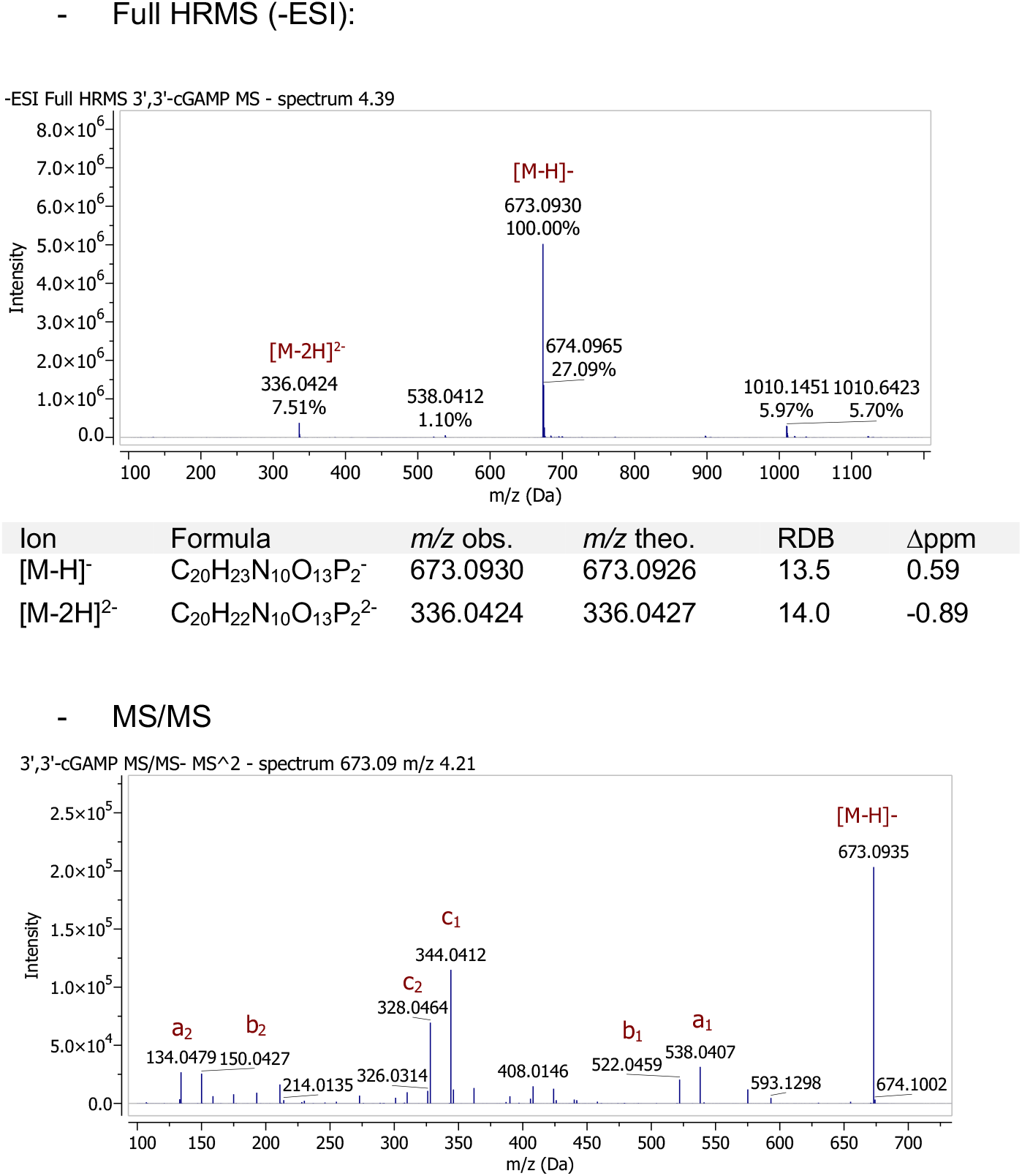
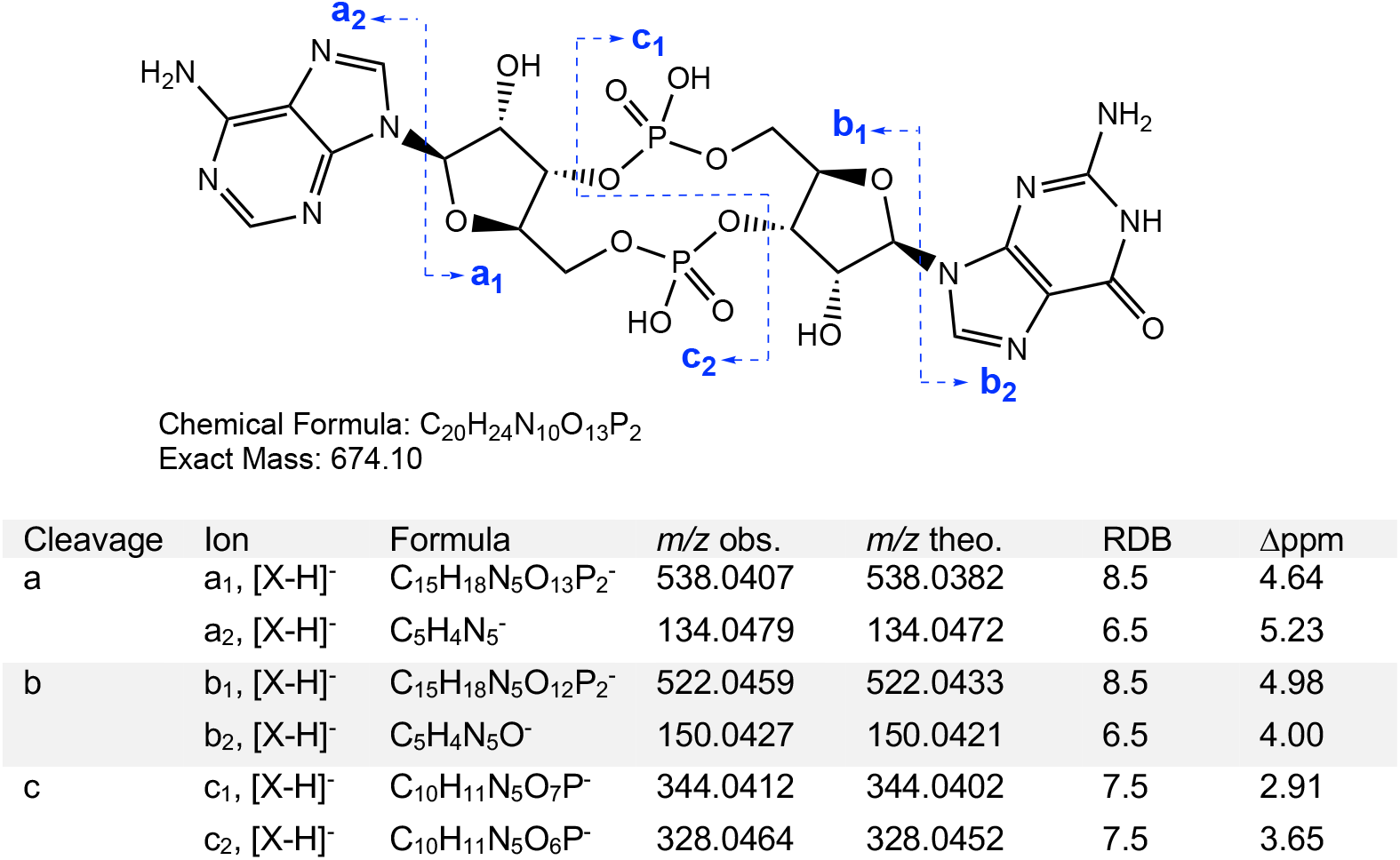

**Characterization of standard 3’,2’-cGAMP:**

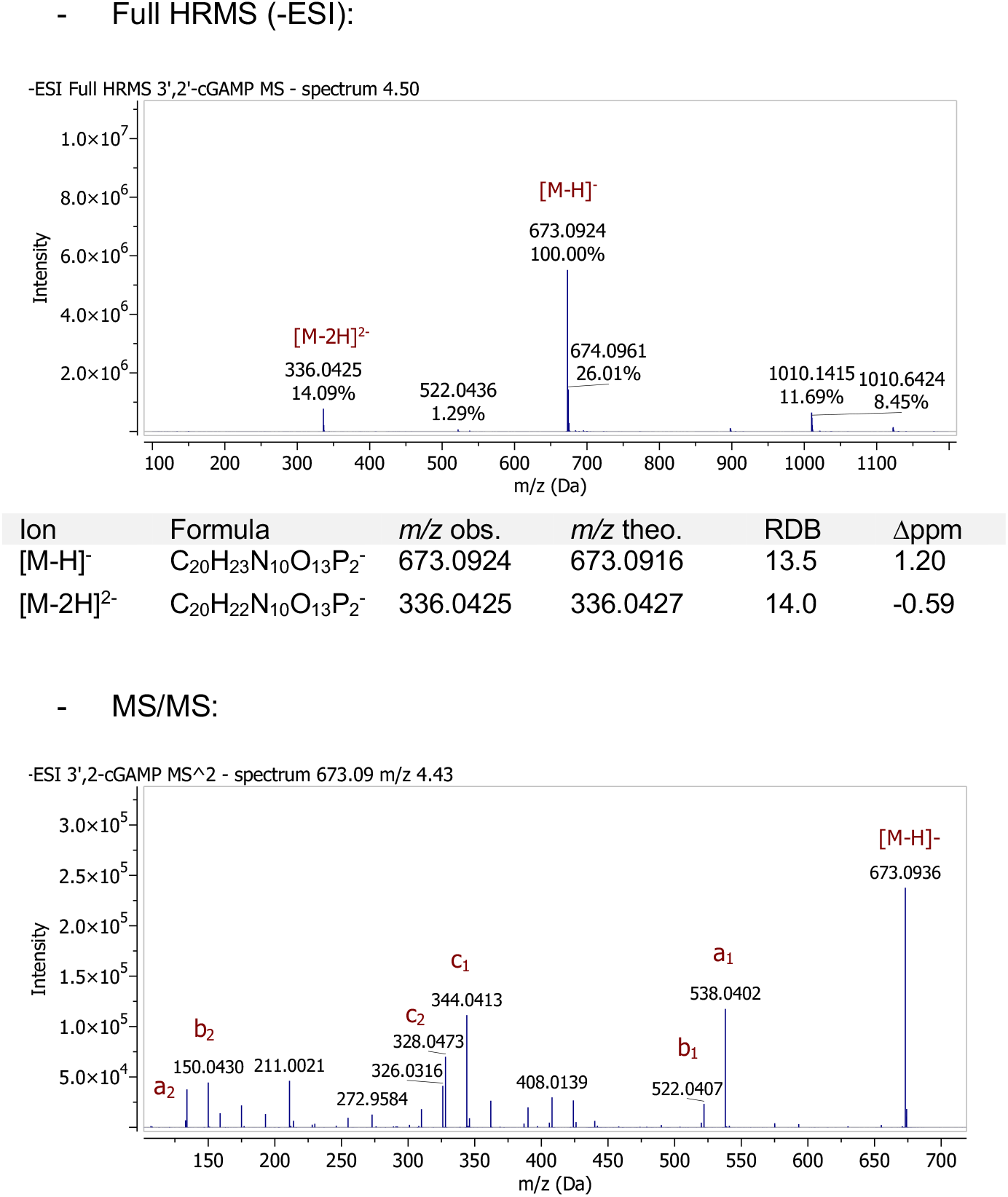
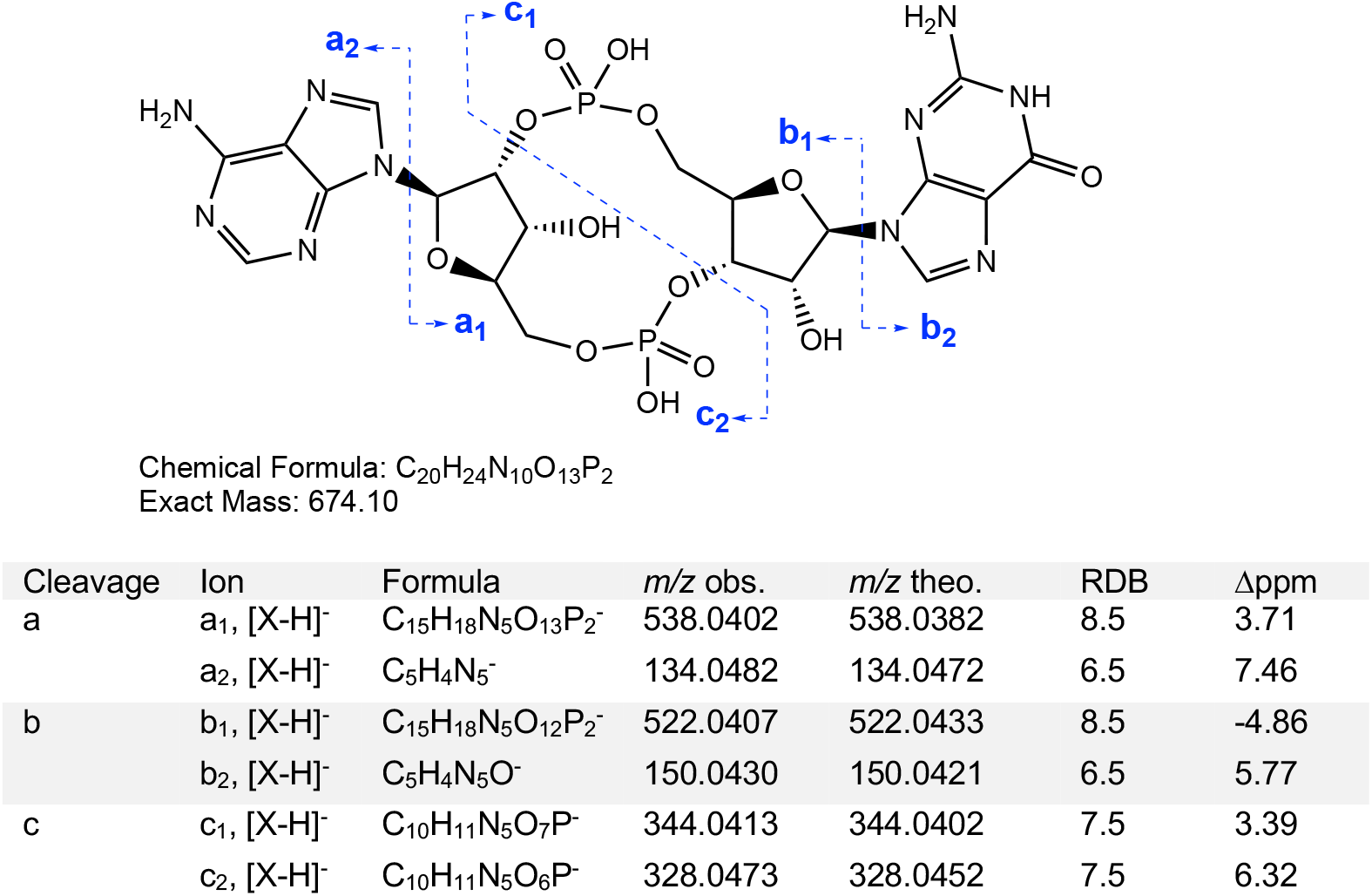

**Characterization of standard 3’,3’-cyclic-dAMP:**

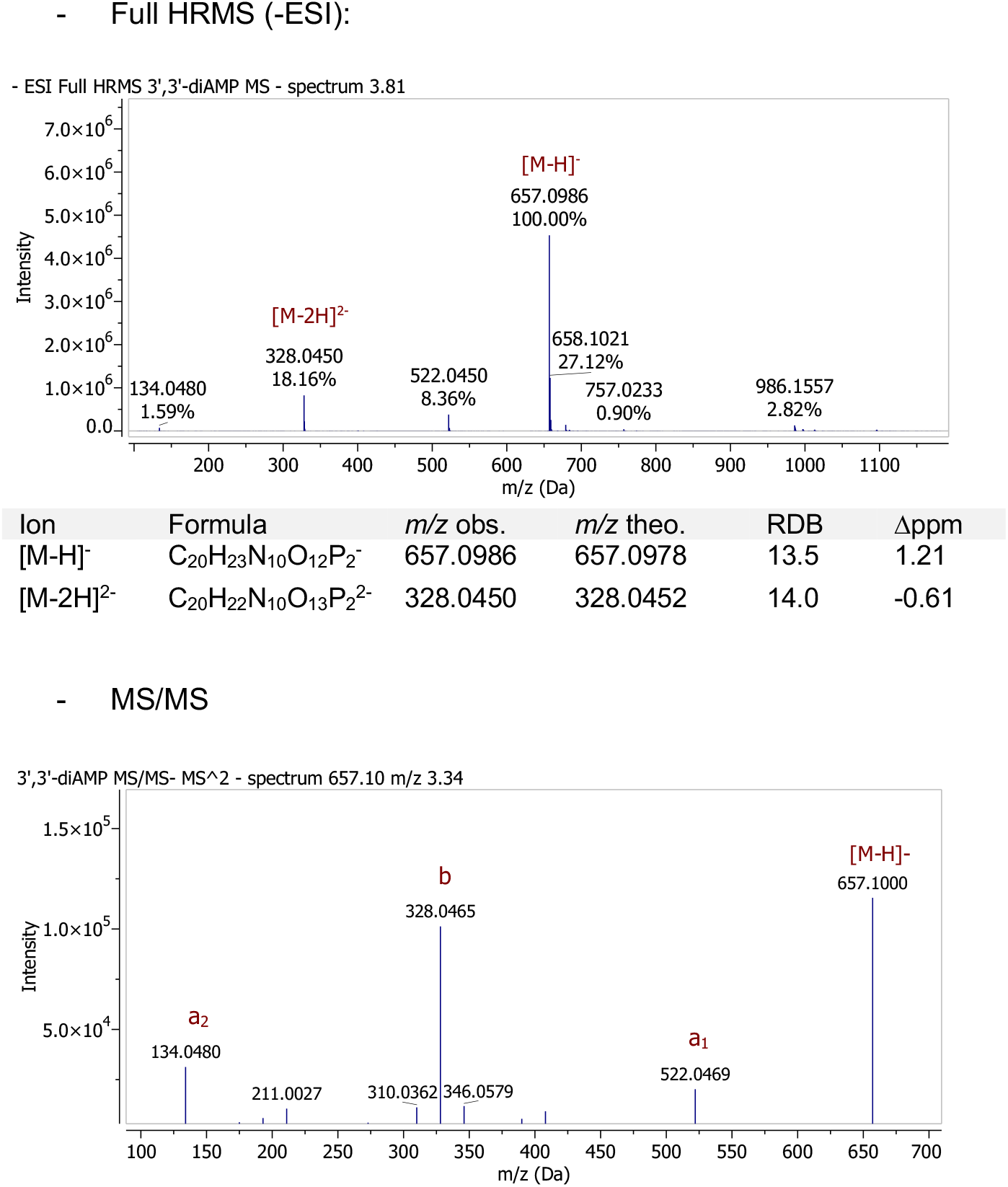
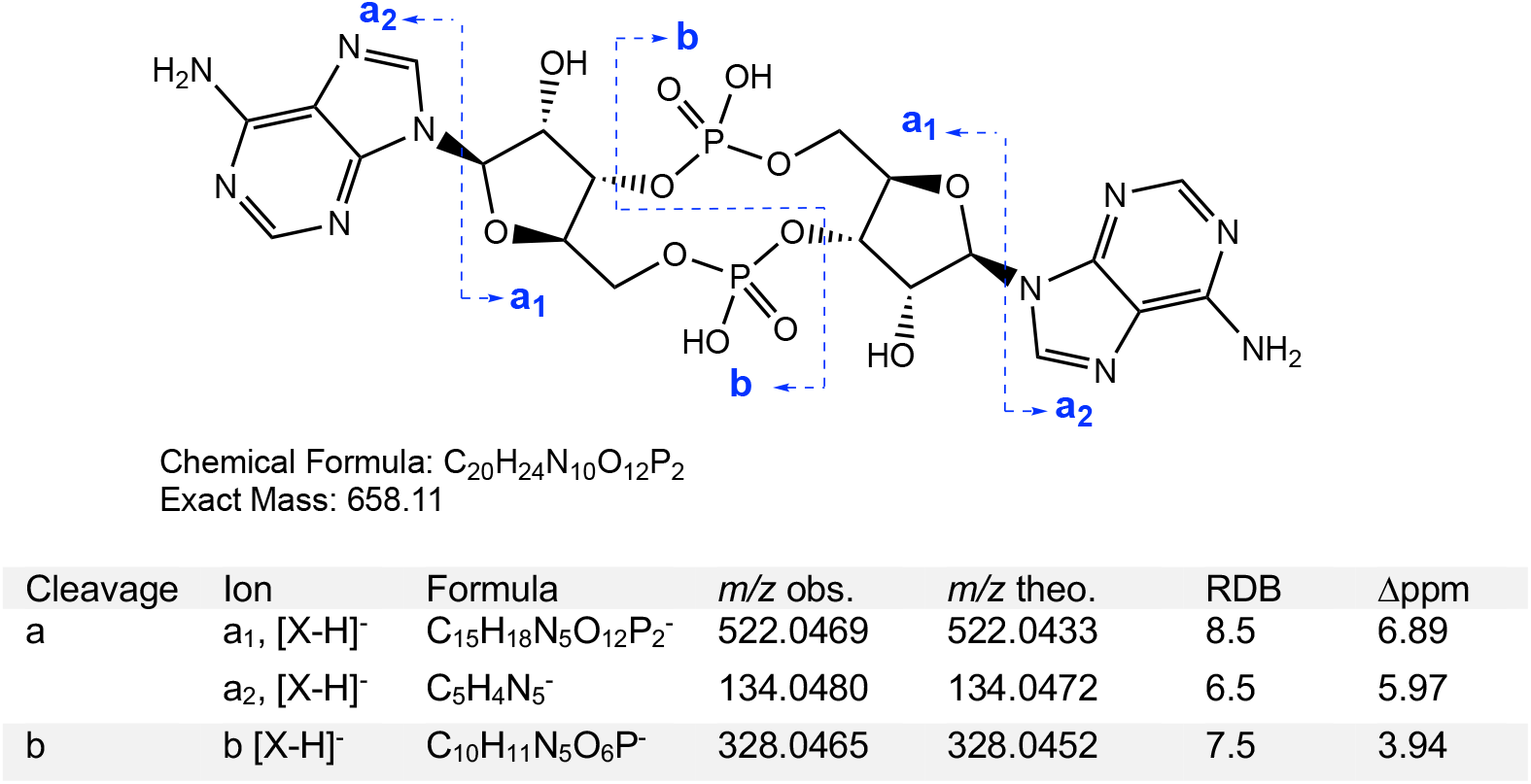

**Characterization of standard 3’,3’-cyclic-dGMP:**

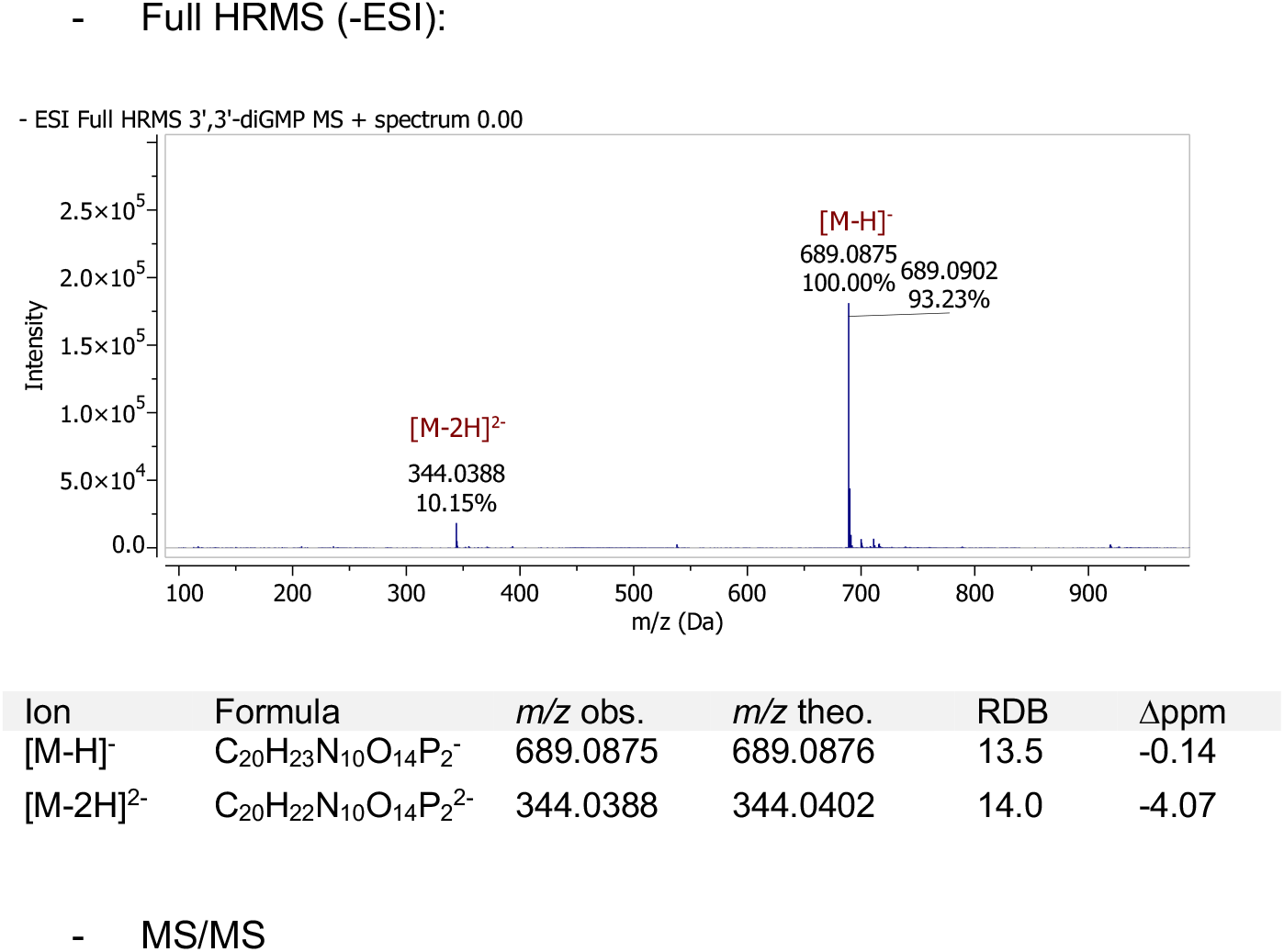
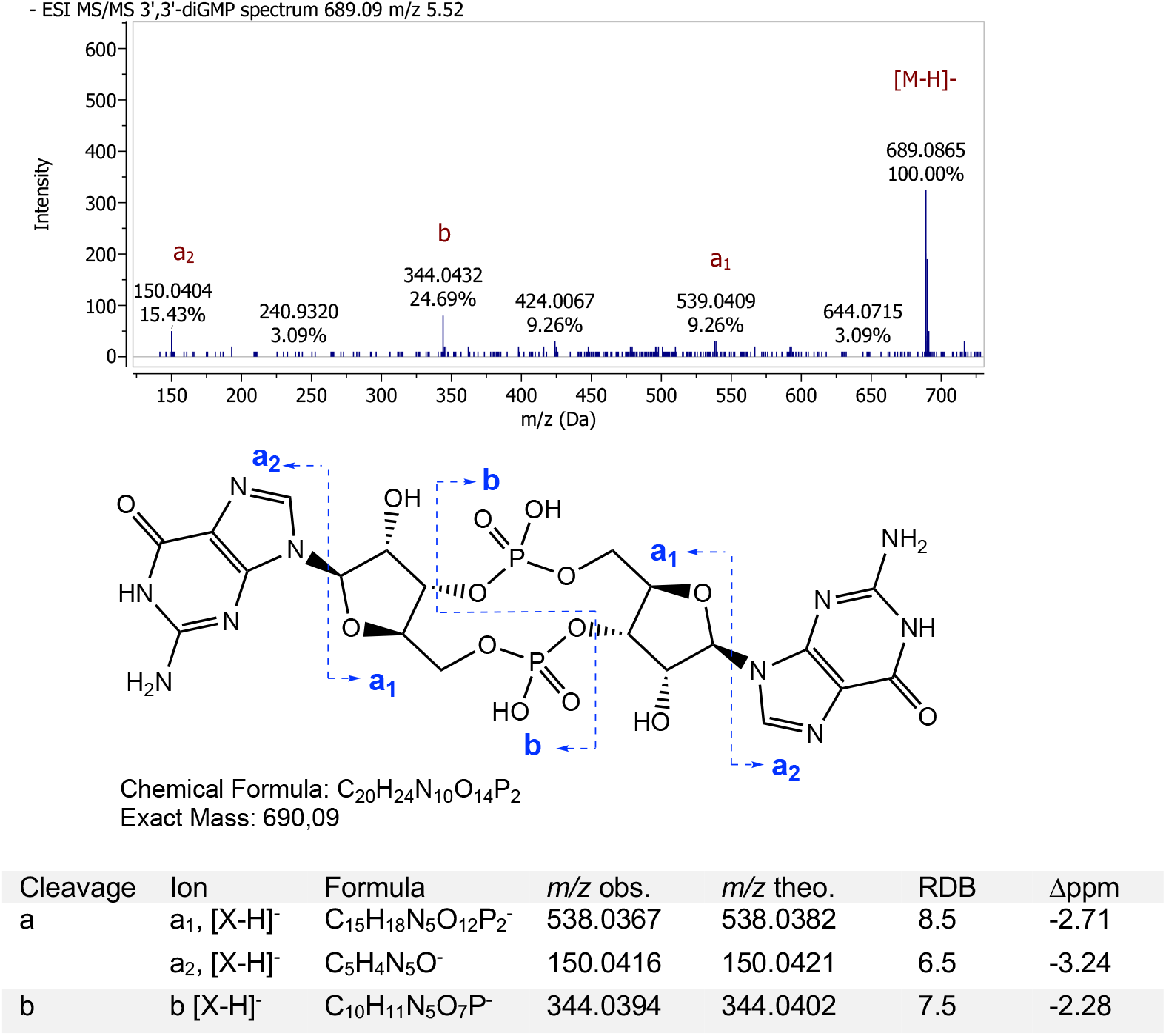

**Analysis of WT enzyme product:**

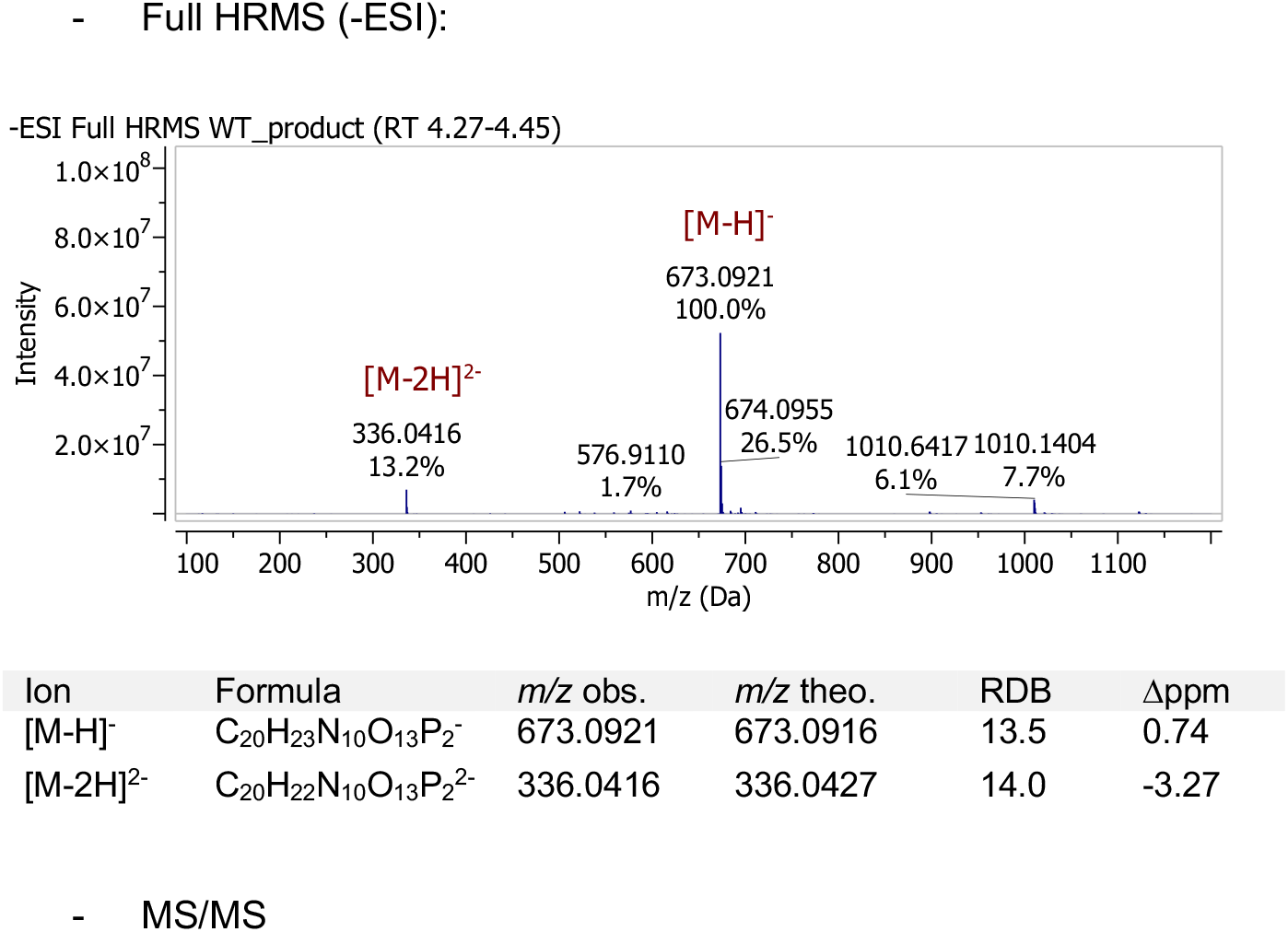
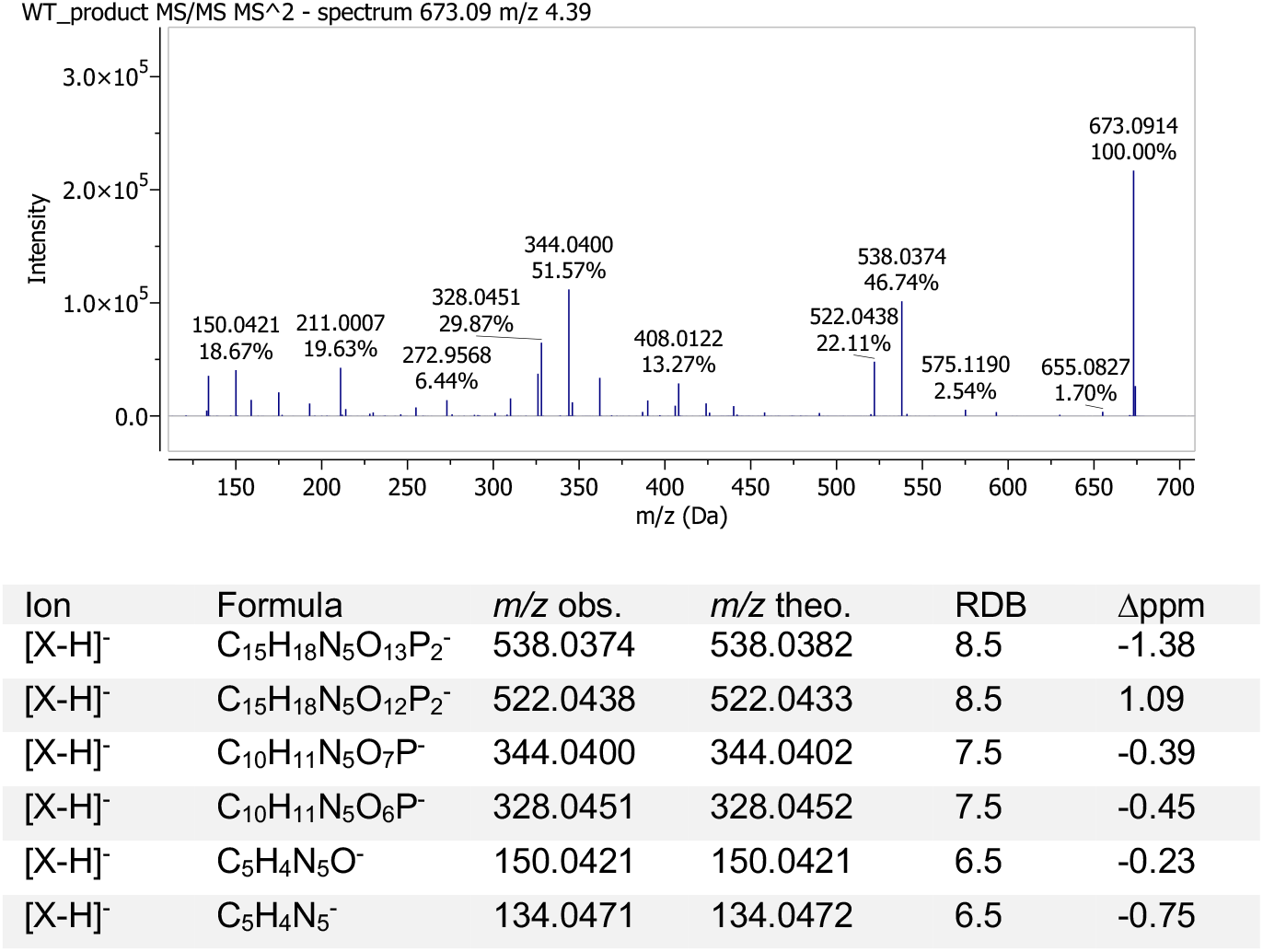

##### POSITIVE ION MODE ACQUISITION

**Characterization of standard 3’,2’-cGAMP:**

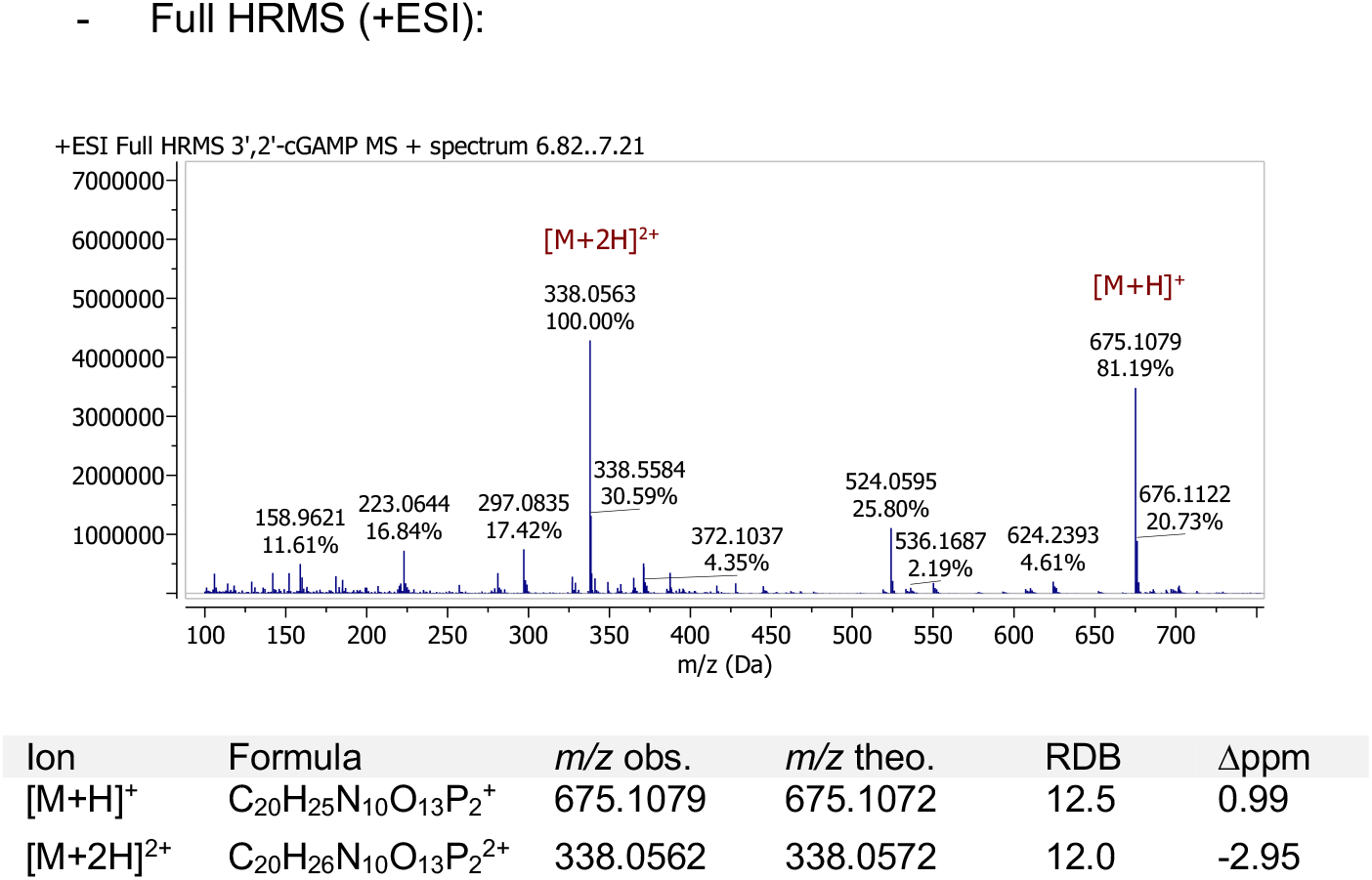

**Characterization of standard 3’,3’-cGAMP:**

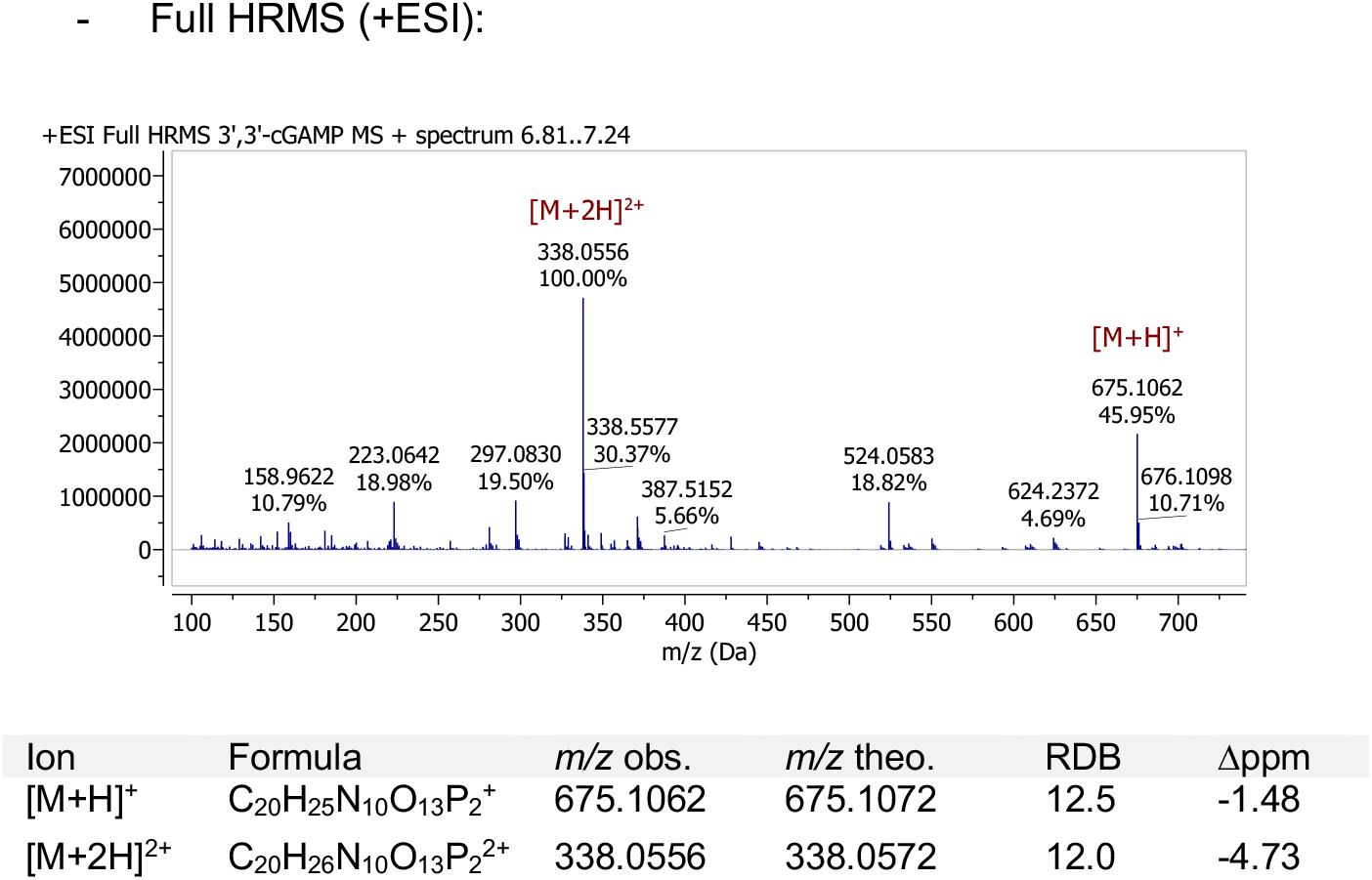

**Characterization of standard 3’,3’-cyclic-dAMP:**

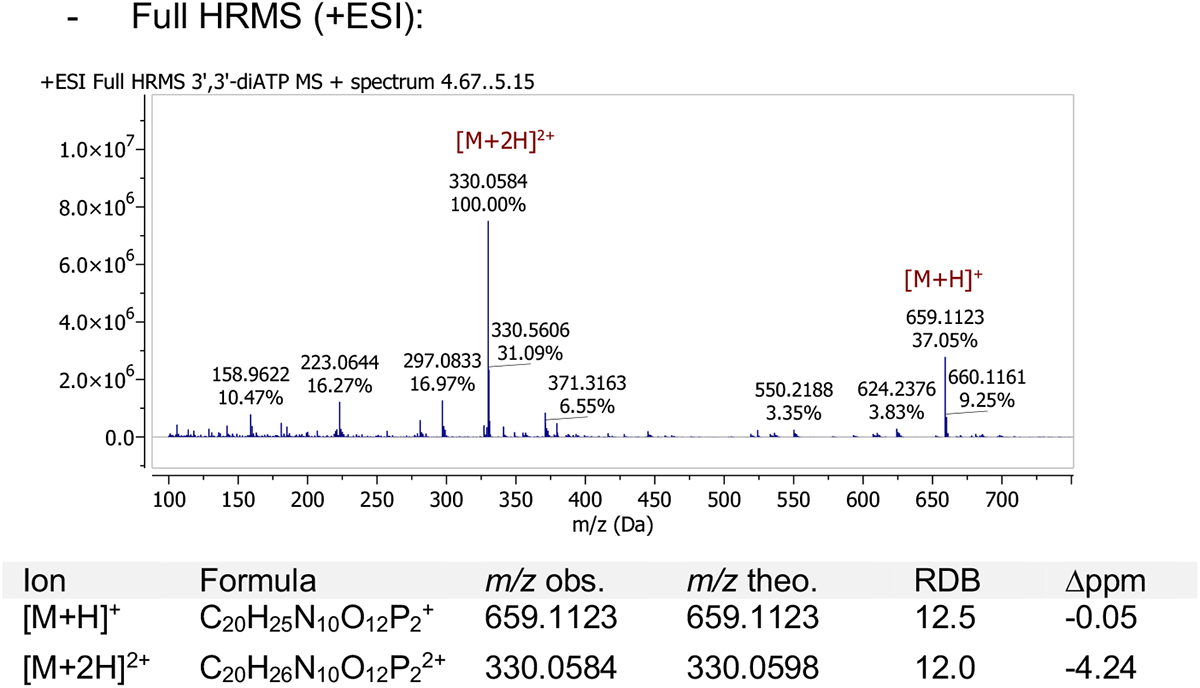

**Characterization of standard 3’,3’-cyclic-dAMP:**

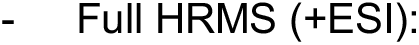
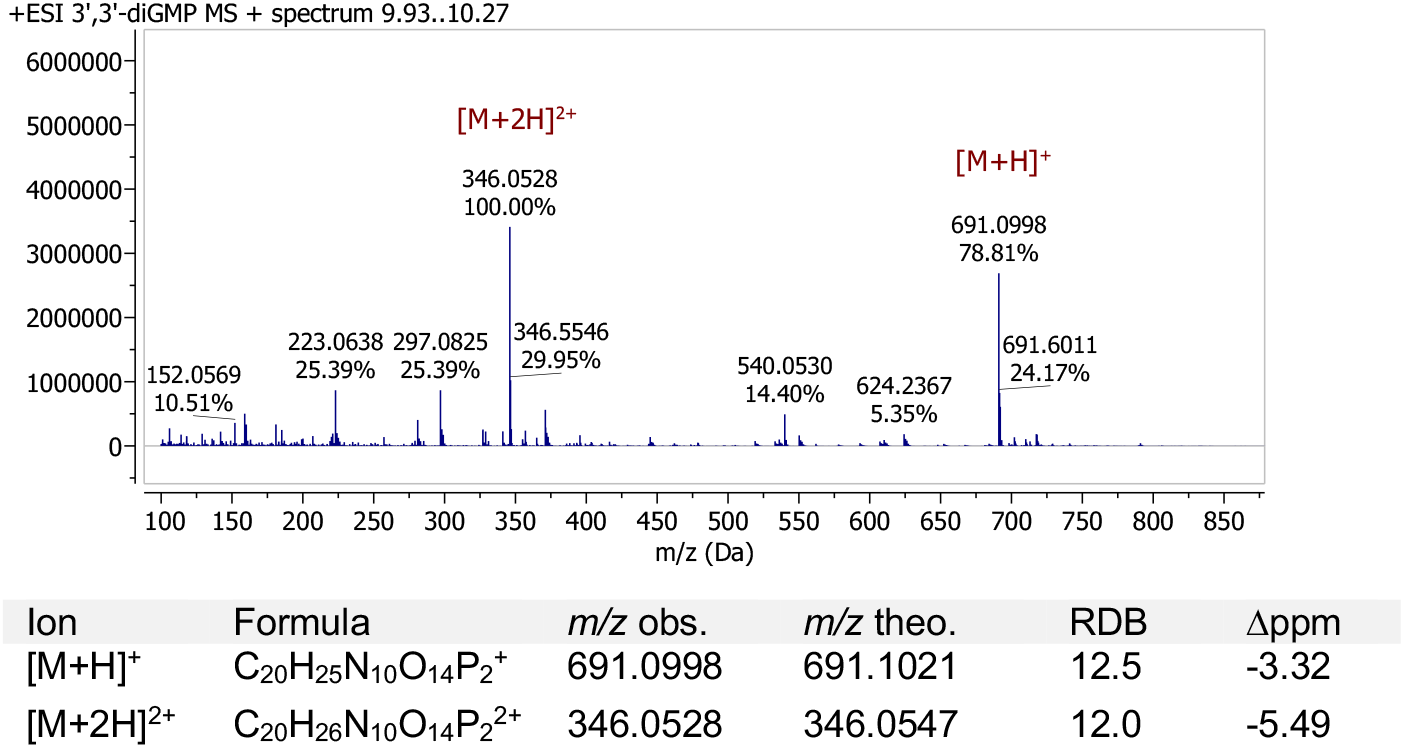

**Analysis of WT enzyme product:**

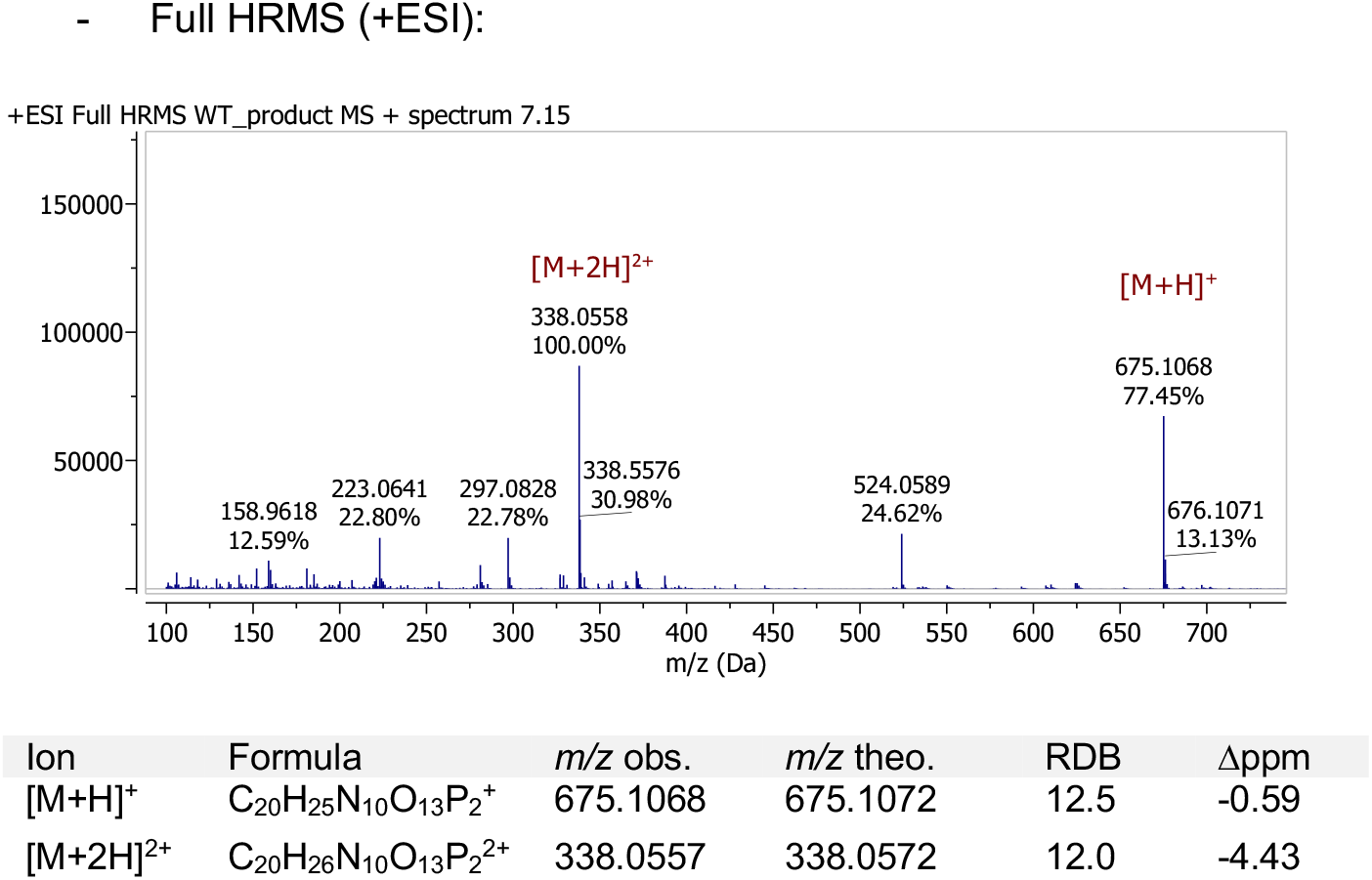

### Figs. S1–S7

**Figure S1.**
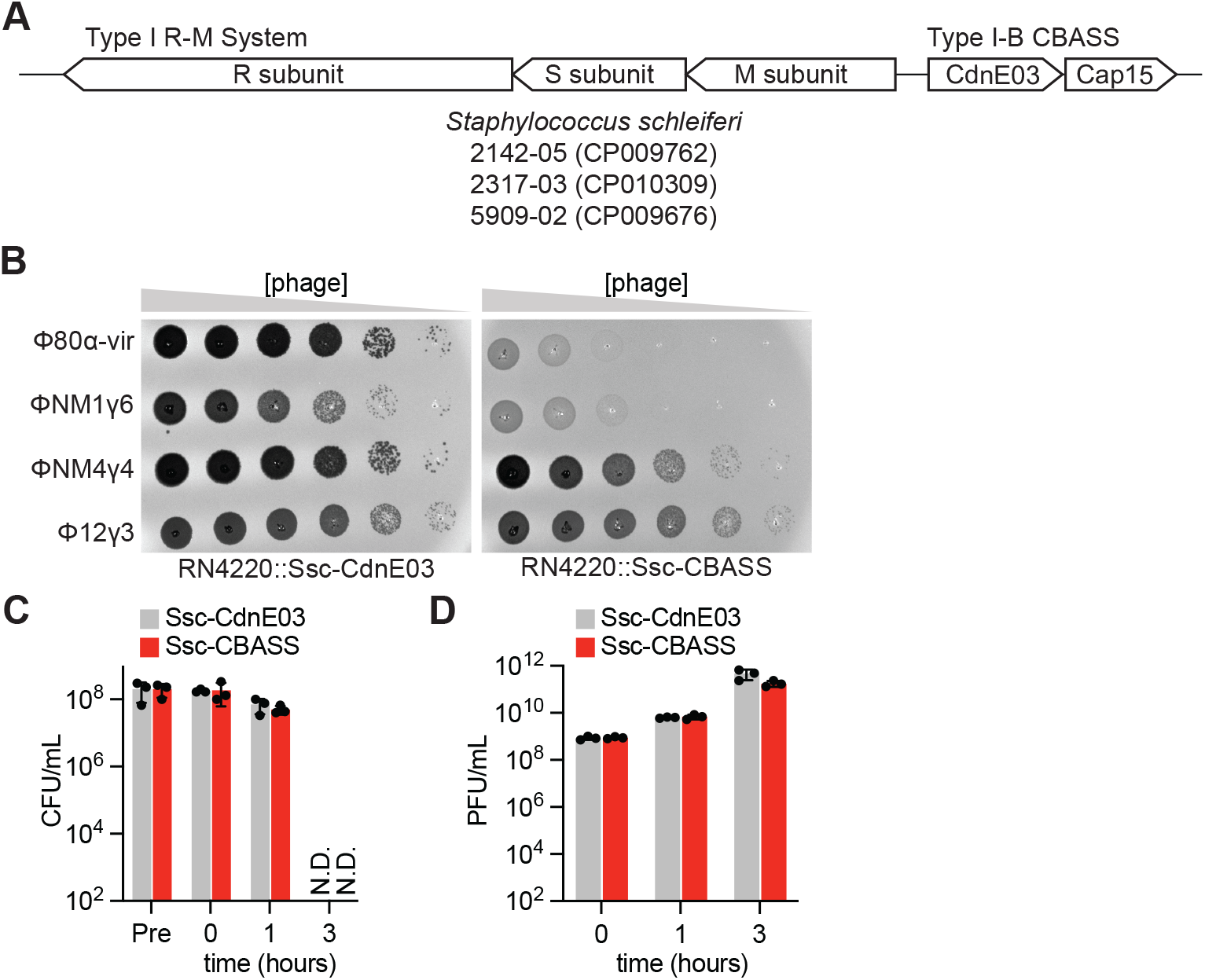
Type I-B CBASS immunity in staphylococci. **(A)** Schematic of the type I-B CBASS operon present in *Staphylococcus schleiferi* (*Ssc*) 2142-05, 2317-03, and 5909-02 genomes, flanked by a type I restriction-modification system. The CBASS operon consists of two genes encoding a cyclase belonging to the E clade, cluster 3 (Ssc-CdnE03) and a two-transmembrane domain-containing effector, Cap15. **(B)** Detection of phage propagation after spotting ten-fold dilutions of the lytic DNA phages, Φ80α-vir, ΦNM1γ6, ΦNM4γ4, and Φ12γ3 onto lawns of *S. aureus* RN4220 harboring either an incomplete (Ssc-CdnE03 alone) or intact Ssc-CBASS operon integrated in its genome. **(C)** Enumeration of colony-forming units (CFU) from cultures harboring Ssc-CdnE03 alone or Ssc-CBASS immediately before infection (Pre), after initial absorption of the phage (0 h), after one lytic cycle (1 h), and after complete culture lysis (3 h) by ΦNM4γ4 at MOI 5. Mean ± SEM of three biological replicates is reported. **(D)** Same as **(C)** but enumerating of plaque-forming units (PFU). Mean ± SEM of three biological replicates is reported.

**Figure S2.**
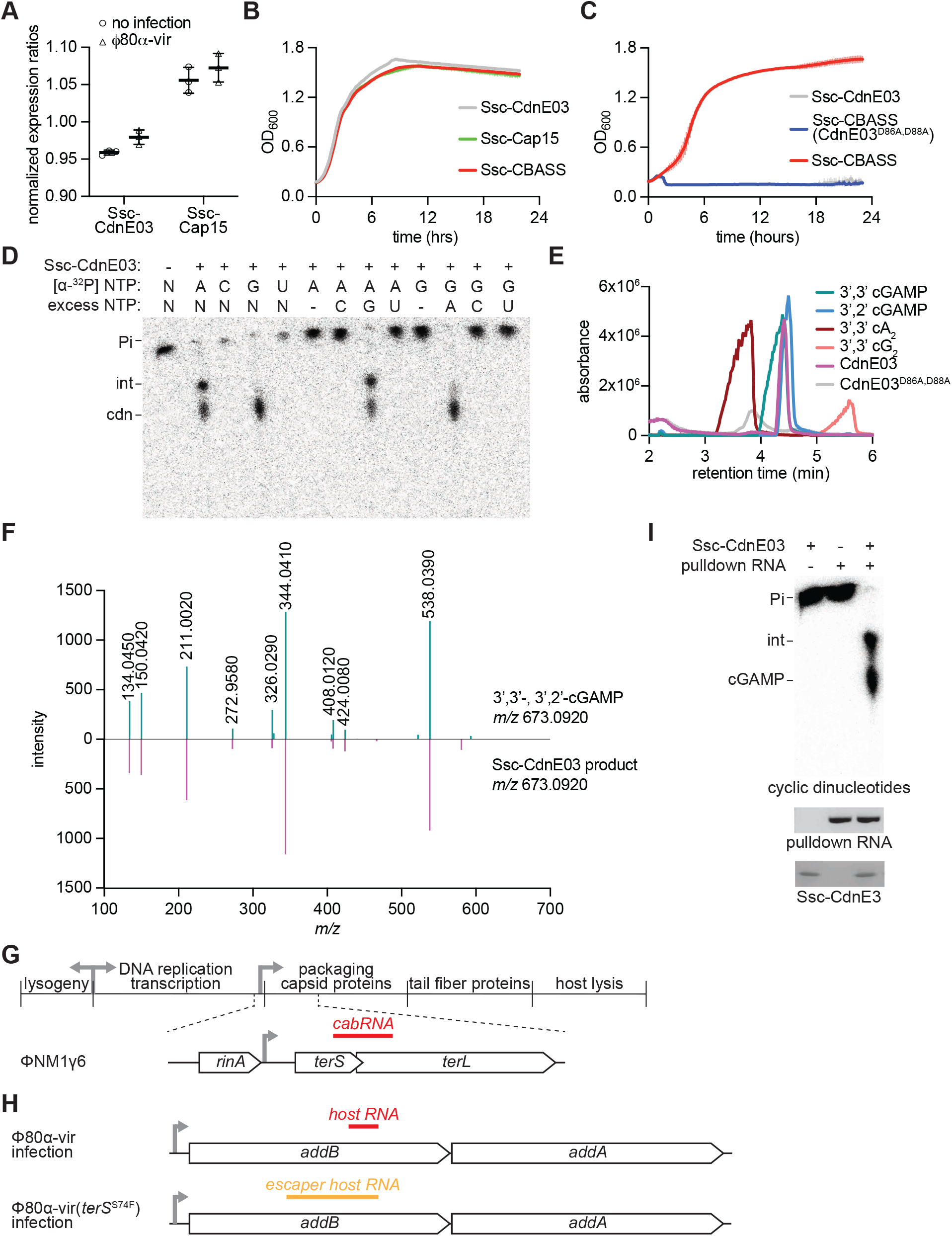
Regulation of Ssc-CBASS operon. **(A)** Expression of the Ssc-CdnE03 and Ssc-Cap15 effector genes during log-phase growth of *S. aureus* RN4220::Ssc*-*CBASS in the presence or absence of infection by Φ80α-vir measured by RT-qPCR. For each condition, expression ratios were determined by normalizing Cq values for Ssc-CBASS genes to Cq values for the housekeeping gene *glcC*. The mean of three biological replicates ± SEM is reported. **(B)** Growth of staphylococci harboring an over-expression plasmid containing either the Ssc-CdnE03 alone, Ssc-Cap15 alone, or the intact Ssc-CBASS operon under the transcriptional control of a P-spac promoter, measured by optical density at 600 nm after the addition of IPTG. The mean of three biological replicates ± SD is reported. **(C)** Growth of staphylococci harboring either an incomplete (Ssc-CdnE03 alone) or intact Ssc-CBASS operon, with either wild-type or D86A,D88A mutant Ssc-CdnE03, measured by optical density at 600 nm after the addition of Φ80α-vir at a MOI of 1. The mean of three biological replicates ± SD is reported. **(D)** Thin-layer chromatography analysis of Ssc-CdnE03 products in the presence of total RNA from *S. aureus* RN4220 after Φ80ɑ-vir infection, using different radiolableled nucleotides to investigate the nucleotide composition of the enzymatic product. Pi, free phosphates; int, intermediate cyclase product; cdn, cyclic dinucleotide. **(E)** TIC of the reaction products of wild-type Ssc-CdnE03. The peak at retention time 4.36 minutes coincides with the retention time of the isomers 3’,2’-cGAMP and 3’,3’-cGAMP. This peak is not present in the reaction products of the active site mutant cyclase, D86A-D88A. **(F)** Comparison of averaged MS/MS spectra of the reaction products of wild-type Ssc-CdnE03 (purple spectrum) and 3’,2’-cGAMP and 3’,3’-cGAMP (green spectrum). The most abundant ions are present in both samples (see Supplementary Text for a complete MS analysis). **(G)** Diagram of the ϕ80α-vir genome showing the localization of the cabRNA sequence. **(H)** Diagram showing the localization of the host RNA and escaper host RNA in the *addAB* operon of *S. aureus* RN4220. **(I)** Thin-layer chromatography analysis of Ssc-CdnE03 reaction products in the presence of cabRNA isolated from a pulldown assay. An agarose gel stained with ethidium bromide (middle) and SDS-PAGE stained with Coomassie blue (bottom) are shown as loading controls. Pi, free phosphates; int, intermediate cyclase product.

**Figure S3.**
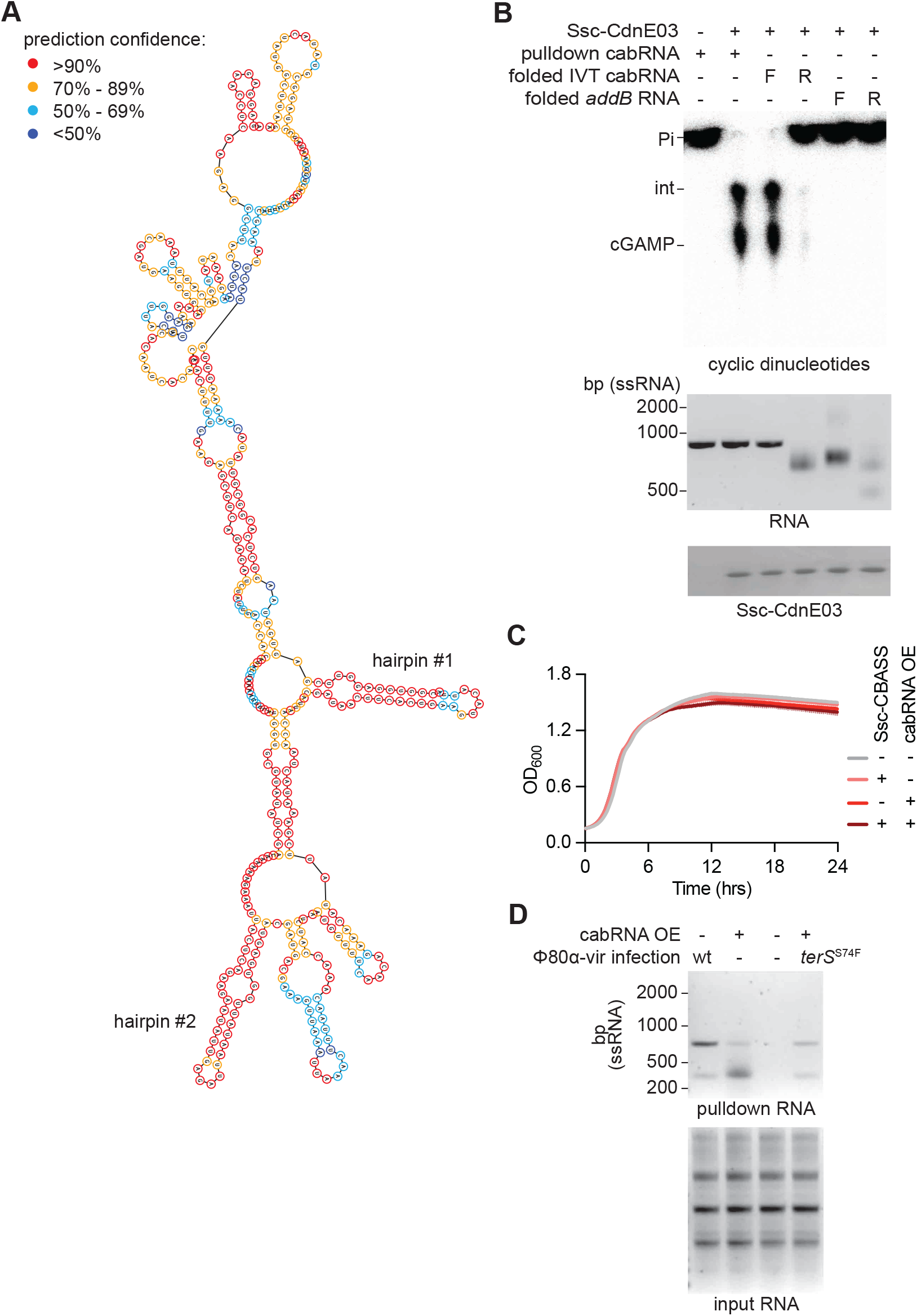
Analysis of cabRNA activation of Ssc-CdnE03. **(A)** Predicted structure of activating RNA. Secondary structure was predicted using ViennaRNA software. Model confidence for each nucleotide is shown as different colors: >=90% (red), 70-89% (orange), 50-60% (light blue), <50% (dark blue). **(B)** Thin-layer chromatography analysis of Ssc-CdnE03 reaction products in the presence of the cabRNA isolated from a pulldown assay, or the sense or antisense strands of *in vitro* transcribed (IVT) cabRNA and host RNA. All IVT RNA was subjected to heat refolding (see Methods). An agarose gel stained with ethidium bromide (middle) and SDS-PAGE stained with Coomassie blue (bottom) are shown as loading controls. Pi, free phosphates; int, intermediate cyclase product. **(C)** Growth of staphylococci harboring either an incomplete (Ssc-CdnE03 alone, “-“) or intact Ssc-CBASS (“+”) operon and an empty vector (“-“) or a plasmid encoding cabRNA (“+”) under the control of an ATc inducible promoter measured by optical density at 600 nm. The mean of three biological replicates ± SD is reported. **(D)** Agarose gel electrophoresis of the input and output RNA obtained after incubation of Ssc-CdnE03 with total RNA extracted from cells infected with Φ80ɑ-vir or Φ80ɑ-vir(*terS*^S74F^) phage, in the presence or absence of cabRNA plasmid overexpression.

**Figure S4.**
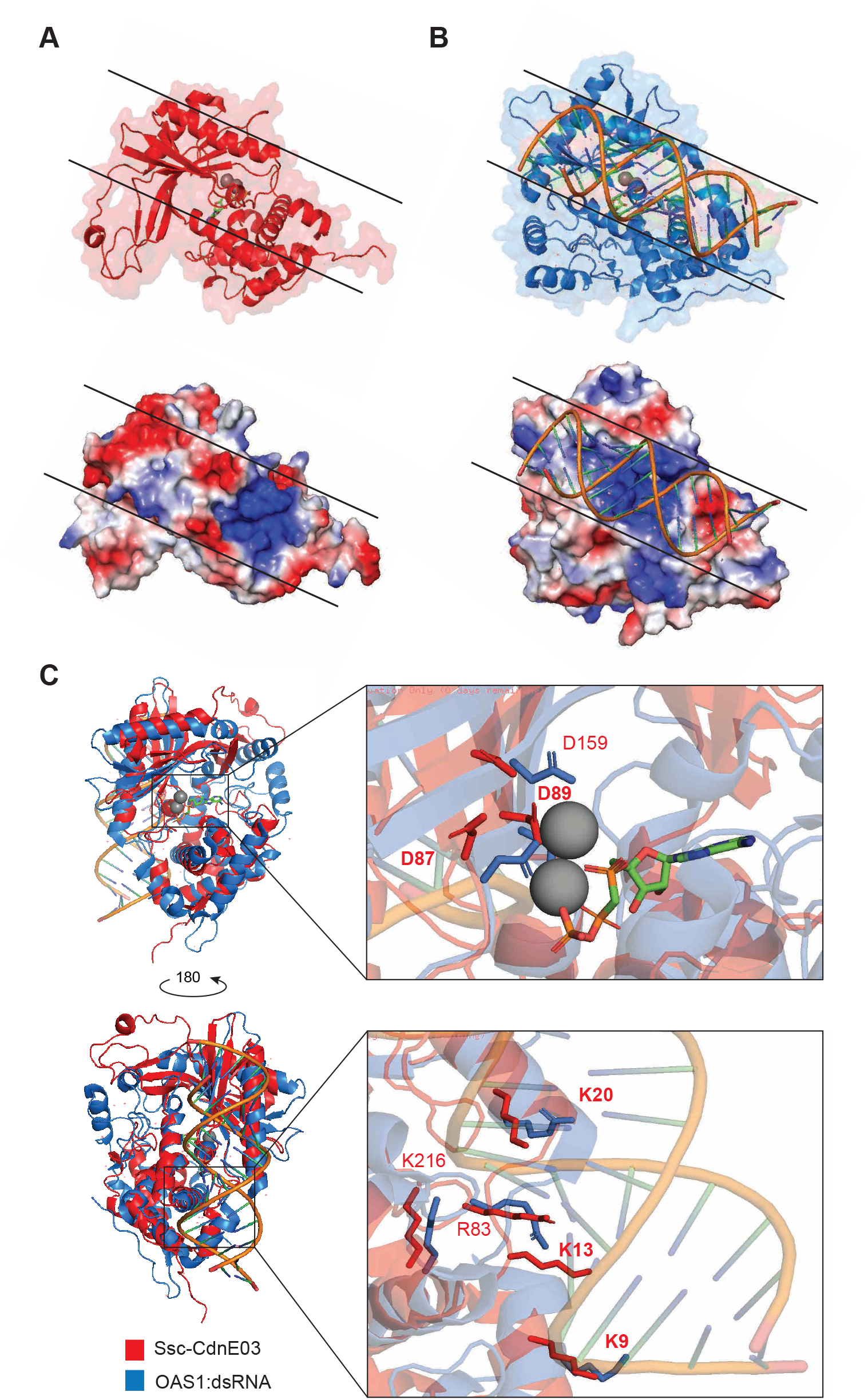
Structural analysis of Ssc-CdnE03. **(A)** AlphaFold rank #1 model of Ssc-CdnE03 (top) with surface electrostatics shown (bottom). Black lines define a conserved primary dsRNA-binding surface present in porcine OAS1. **(B)** Crystal structure of porcine OAS1 bound to dsRNA (PDB: 4RWO) (top) with surface electrostatics shown (bottom). Black lines define the dsRNA-binding surface. **(C)** Structural alignment of Ssc-CdnE03 (red) and crystal structure of porcine OAS1:dsRNA (PDB: 4RWO) (blue) with zoomed-in cutaways highlighting conservation of the active site (top inset) and positively charged residues within the ligand binding surface (bottom inset).

**Figure S5.**
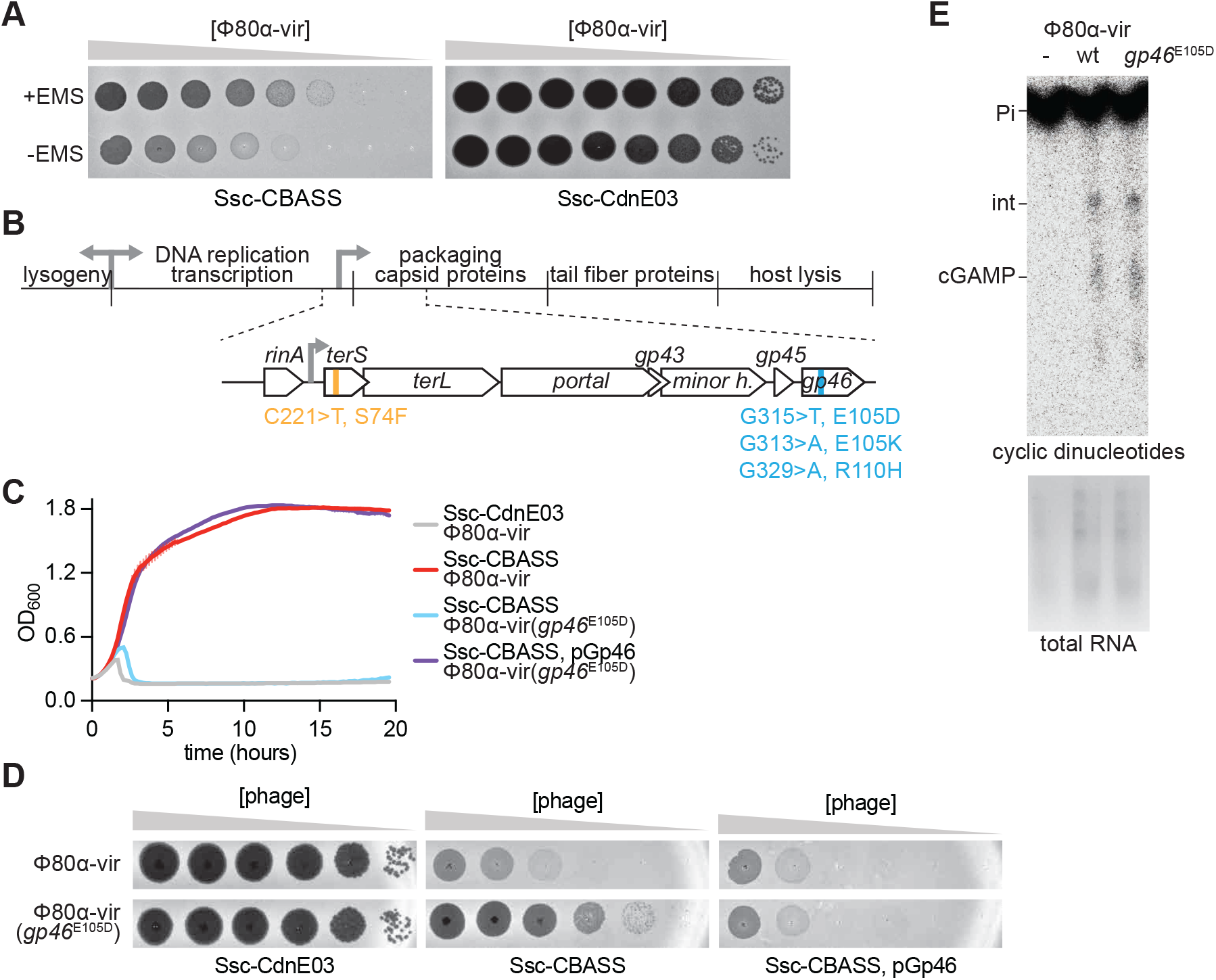
Isolation and characterization of CBASS escapers. **(A)** An overnight culture of *S. aureus* RN4220 was diluted and outgrown to early log-phase, at which time Φ80ɑ-vir at an MOI of 1 was added. Just before the first burst (∼30 min), 1% ethyl methanesulfonate (EMS) was added to generate mutations. Infections in the presence of EMS were allowed to proceed at 37°C for 4 hours to allow phage to propagate and lyse the culture. Culture supernatants were collected, serially diluted, and spotted on a lawn of *S. aureus* RN4220::Ssc-CBASS or Ssc-CdnE03. A control experiment without the addition of the EMS mutagen is shown as control. **(B)** Diagram of ϕ80α-vir genome with localization of four unique escaper mutations identified in *terS* or *gp46*. **(C)** Growth of staphylococci harboring either an incomplete (Ssc-CdnE03 alone) or intact Ssc-CBASS operon measured by optical density at 600 nm after the addition of Φ80α-vir or ϕ80α-vir(*gp46*^E105D^) at MOI 1. The mean of three biological replicates ± SD is reported. **(D)** Detection of phage propagation after spotting ten-fold dilutions of Φ80α-vir or ϕ80α-vir(*gp46*^E105D^) onto lawns of *S. aureus* RN4220 harboring either an incomplete (Ssc-CdnE03 alone) or intact Ssc-CBASS operon, the latter in the presence or absence of Gp46 overexpression using plasmid pGp46. **(E)** Thin-layer chromatography analysis of Ssc-CdnE03 reaction products in the presence of total RNA extracted from uninfected staphylococci or cells infected with wild-type or *gp46*^E105D^ Φ80ɑ-vir. Agarose gel electrophoresis of RNA samples is displayed as a loading control.

**Figure S6.**
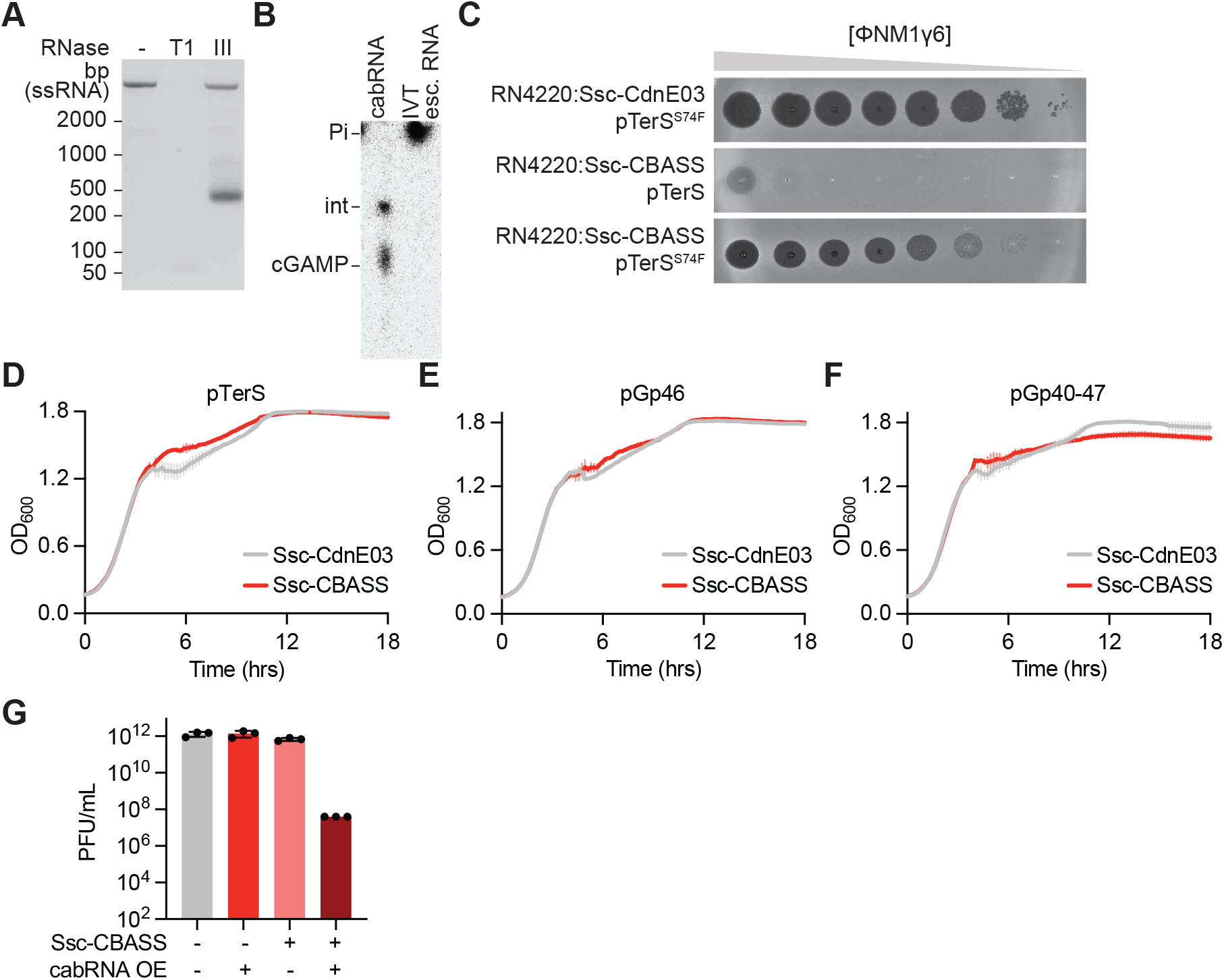
Mechanism of escape mediated by the *terS* mutation. **(A)** Agarose gel electrophoresis of the escaper RNA generated during infection with ϕ80α-vir(*terS*^S74F^) isolated from a pulldown assay, treated with RNase T1 or III. **(B)** Thin-layer chromatography analysis of Ssc-CdnE03 reaction products in the presence of cabRNA or *in vitro* transcribed escaper RNA. **(C)** Wild-type ΦNM1γ6 was propagated in liquid cultures of *S. aureus* RN4220::Ssc-CBASS harboring a plasmid-borne *terS* gene, wild-type or S74F. Culture supernatants were collected and serial dilutions were spotted onto lawns of *S. aureus* RN4220::Ssc*-*CdnE03 or RN4220::Ssc*-*CBASS. **(D)** Growth of staphylococci harboring pTerS, providing IPTG-inducible expression of the Φ80α-vir TerS protein, measured by optical density at 600 nm after the addition of the inducer. The mean of three biological replicates ± SD is reported. **(E)** Same as **(D)** but using pGp46 plasmid, providing IPTG-inducible expression of the Φ80α-vir Gp46 protein. **(F)** Same as **(D)** but using pGp40-47 plasmid, providing IPTG-inducible expression of the complete Φ80α-vir viral capsid. **(G)** Enumeration of plaque-forming units (PFU) from cultures harboring Ssc-CdnE03 alone (“-“) or Ssc-CBASS (“+”) and either an empty vector (“-“) or a plasmid with cabRNA (“+”) under the control of an aTc-inducible promoter.

**Figure S7.**
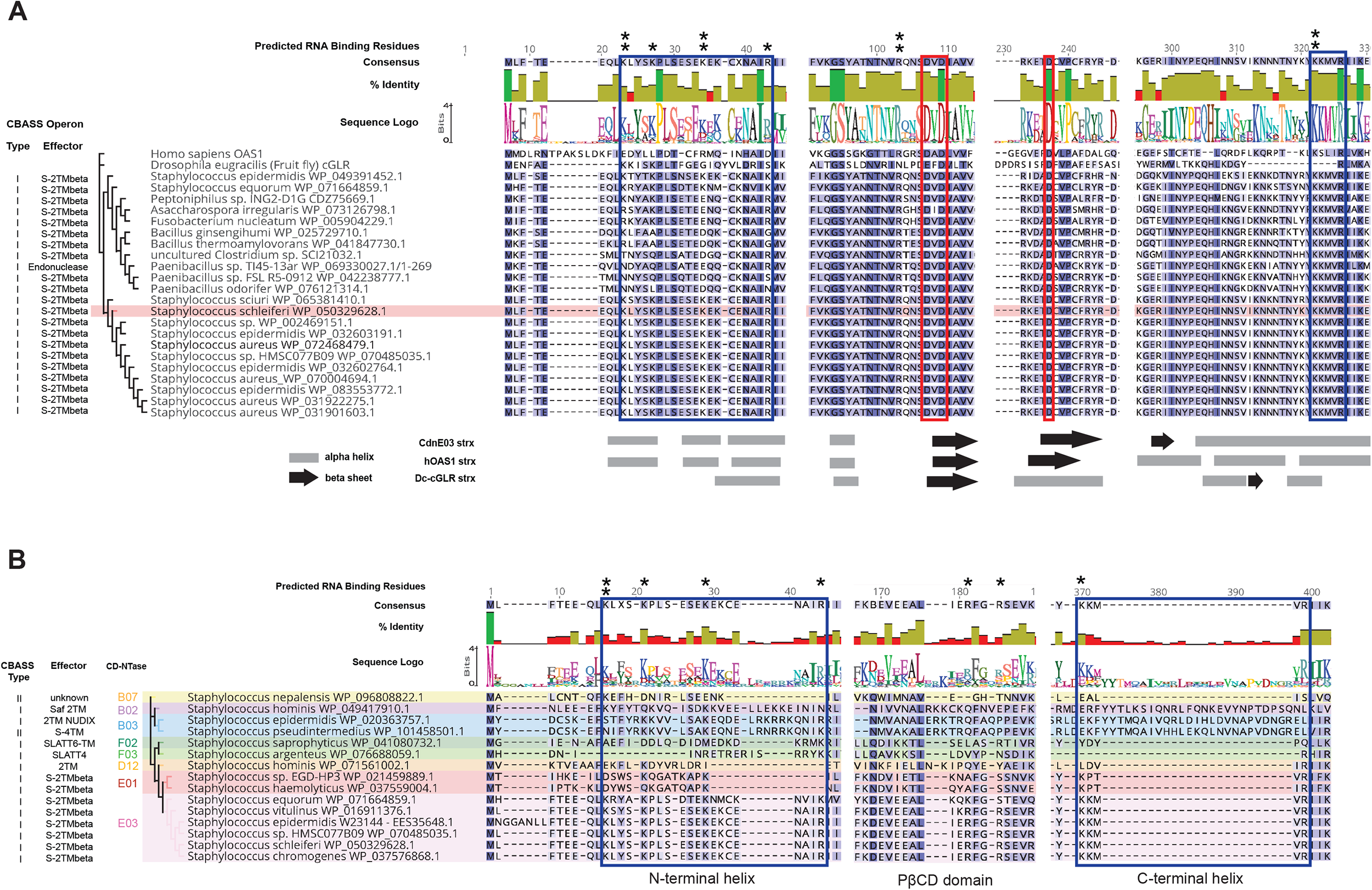
Sequence analysis of CdnE03s. **(A)** Alignment of porcine OAS1, *D. erecta* cGLR, and bacterial CdnE03s and **(B)** Alignment of CD-NTases from staphylococcal species. The EhD[X50-90]D catalytic triad is highlighted with a red outline and the residues that make up the ligand binding site are highlighted with a blue outline. Predicted basic ligand binding residues selected for mutational analysis are denoted with black stars (*= conserved in bacteria, **= conserved in bacteria and OAS1).

